# Spatial expression pattern of *ZNF391* gene in the brains of patients with schizophrenia, bipolar disorders or major depressive disorder identifies new cross-disorder biotypes: A trans-diagnostic, top-down approach

**DOI:** 10.1101/768515

**Authors:** Hongyan Ren, Yajing Meng, Yamin Zhang, Qiang Wang, Wei Deng, Xiaohong Ma, Liansheng Zhao, Xiaojing Li, Yingcheng Wang, Pak Sham, Tao Li

## Abstract

**Background:** Given the struggle in the field of psychiatry to realize the precise diagnosis and treatment, it is in an urgent need to redefine psychiatric disorders based on objective biomarkers. The results generated from large psychiatric genomic consortia show us some new vantage points to understand the pathophysiology of psychiatric disorders. Nevertheless, how to relate these captured signals to the more refined disease dimensions has yet to be established.

**Methods:** We chose a top-down, cross-disorder approach by using the summary statistics of GWAS from large psychiatric genomic consortia to build a genomic structural equation model (SEM) for SCZ, BD and MDD to detect their common factor (CF), and to map a potential causal core gene for the CF, followed by the transcriptional prediction of the identified causal gene in our sample and the discovery of new biotypes based on the prediction pattern of the causal gene in the brain. We then characterized the biotypes in the context of their demographic features, cognitive functions and neuroimaging traits.

**Outcomes:** A common factor emerged from a well-fitting genomic SEM of SCZ, BD and MDD (loading 0.42, 0.35 and 0.09 for SCZ, BD and MDD, respectively). One genomic region in chromosome 6 was implicated in the genetic make-up of the common factor, with fine-mapping analysis marking *ZNF391* as a potential causal core gene (posterior possibility = 0.96). Gene expression inference analysis identified eight brain regions showing different expression levels of *ZNF391* between patients and controls, with three biotypes arising from clustering patients based on their expression pattern of *ZNF391* in the brain. The three biotypes performed significantly differently in working memory (P_DMS_TC_ = 0.015, P_DMS_TC_A_ = 0.0318, P_DMS_t0D_ = 0.015) and demonstrated different gray matter volumes in right inferior frontal orbital gyrus (RIFOG) in the same order as working memory (biotype 3 > biotype 2 > biotype 1, P_RIFOG_ = 0.0027). Using *ZNF391* as instrumental variable (IV), a partial casual path could be linked from RIFOG to working memory (**β**_RIFOG->DMS_TC0D_ = 4.95, P = 0.0056; **β** _RIFOG->DMS_TC_ = 2.53, P = 0.059; **β**_RIFOG->DMS_TC_A_ = 2.57, P = 0.056).

**Interpretation:** The general predisposition to several psychiatric disorders may be influenced by variations of ZNF391, through its effects on right inferior frontal orbital gyrus and working memory. This illustrates the potential of a trans-diagnostic, top-down approach in understanding the commonality of psychiatric disorders.

**Evidence before this study:** The results from recent cross-disorder genome-wide association studies (GWAS)using large samples indicate that there is notable genetic overlapping between psychiatric disorders. However, the structural relationship of these disorders at the genomic level and the details of refined disease dimensions affected by the associated loci in a cross-disorder pattern remains unknown. We searched the published studies (up to Sep 7, 2019) in PubMed using the combination of the following keywords “((cross disorder) OR (schizophrenia AND bipolar disorder AND major depressive disorder) AND (genome AND structural equation) AND (cognition OR imaging))”, no published study was found. We then removed the term “structural equation”, 23 original studies were found. To the best of our knowledge, none of these studies explored the organized structure between three disorders. Further, of 23 articles we found, the majority of them took an approach of either polygenic risk score (PRS) or candidate gene to test the association with either psychological traits such as loneliness or neuroimaging measures in one (schizophrenia) or two (schizophrenia and bipolar) disorders. Hitherto, no study has been conducted to redefine three disorders based on the biological markers generated from the cross-disorder genomic studies.

**Added value of this study:** Adopting a novel approach of genomic structural equation modelling, we used the latest results of GWAS of three major psychiatric disorders to detect their common factor, further, to identify the loci associated with such as a common factor, and the loci’s transcription consequences in the brain. Propelled by the phenomenon “genes do not read DSM”, we used a cutting-edge clustering algorithm to redefine three disorders based on the cerebral spatial expression pattern of associated core gene. Our study provides another piece of evidence as to the potentials of utilizing the signals arising from large population-scale GWAS to dissect and redefine psychiatric disorders.

**Implications of all the available evidence:** Consistent with previous case-control cross-disorder GWAS, our study suggests that a common factor exists in three major psychiatric disorders and the biological information of core gene associated with the common factor could be used as an objective marker to explain three disorders and their pathophysiology.

## Introduction

Thanks to the concerted efforts from multi-national consortia, remarkable progress has been made in the field of psychiatric genetics ^1-3^. However, translating the findings from large-scale genome-wide association studies (GWAS) to clinical applications remains elusive. The two main reasons for such a dilemma are the phenotypic and genetic heterogeneity in the affected populations, due to the current lack of understanding of the pathogenesis of psychiatric disorders. Both classical family studies and recent genome-wide association/sequencing studies have observed a notable co-occurrence / co-segregation of multiple psychiatric disorders ^4,5^. Such findings imply that a cross-disorder approach could increase statistical power to detect susceptibility genes and provide a fuller picture of the genetic relationships between psychiatric disorders ^6^.

Moreover, a large proportion of the SNPs identified by GWAS so far are located at intergenic/non-coding regions, making direct interpretation of the association signals a difficult task ^7^. However, with the establishment of gene expression data repositories, such as GTEx ^8^ and psychENCODE^9^, it has become possible to link many non-coding SNPs (eQTL) to the expression levels of their target genes. By incorporating the eQTL information and tissue-specific expression data, methods have been developed to enable the prediction, in an independent sample, of gene expression level in a tissue of interest ^10^.

Motivated by these new trends, this study seeks to (1) identify the shared latent factor of schizophrenia (SCZ), bipolar disorder (BD) and major depressive disorder (MDD), by using a novel cross-disorder genomic structural equation model (genomic SEM), (2) evaluate the expression levels of potential core genes associated with such a shared factor in the brain regions, and (3) re-delineate patient groups based on distinctive expression patterns (biotypes), and annotate the new biotypes in terms of scalable traits such as cognitive functions and gray matter volumes (the schematics of the study design is shown in Figure 1).

**Figure 1:**
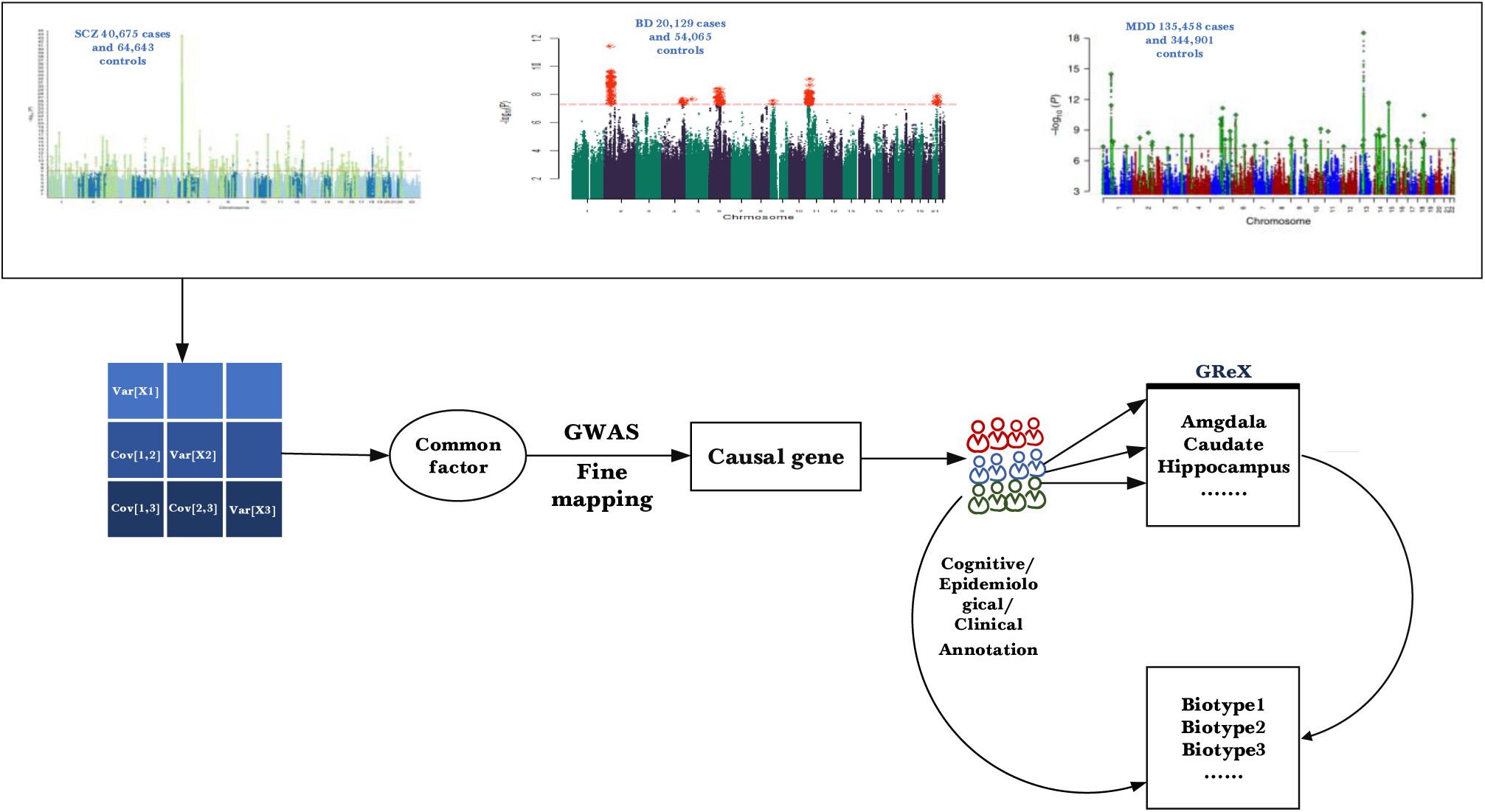
A schematic for data analysis workflow of our study. We started by a genomic SEM modelling of genetic covariance matrix generated from the summary statistics of published GWASs; then we identified the causal gene of the common factor underlying three psychiatric disorders, followed by the imputation of genetically regulated expression (GReX) of the causal gene in the brain. Subsequently, we re-clustered the patient group based on their GReX profile and annotated the clusters in terms of epidemiological, cognitive and neuroimaging indexes

## Methods

### *Genomic SEM and in silico* fine-mapping

We leveraged the summary statistics from large-scale GWAS of three disorders ^11-13^ to model their latent structure using genomic SEM^14^. The methodological details of genomic SEM are described in the appendix (pp 1). Following the emergence of the common factor (CF) underlying the disorders, we implemented a genome-wide scan to map loci conferring effect on CF. In brief, the same SEM model was extended to incorporate SNP effects on each disorder based on GWAS summary statistics, to estimate the effect of SNP on the CF, with corresponding standard error and P-value. The extent to which an SNP’s effect on the three disorders is not mediated through CF is measured by a heterogeneity Q value, with a P-value to indicate its statistical significance. The mapped loci were annotated using FUMA ^15^; the estimation of the heritability explained by SNPs and the genetic correlation analysis between CF and other phenotypes were performed using the LD hub platform ^16^.

To further locate the potential causal core SNPs/genes, we carried out an *in silico* fine-mapping analysis in the mapped region. We chose two algorithms with different theoretical bases to ensure robustness; relevant details of the algorithms (“FINEMAP”^17^and “iRIGS”^18^) are given in the appendix (pp 1).

### Expression inference of ZNF391 in the brain and cluster analysis

The fine-mapping above identified zinc finger protein 391 (*ZNF391)* as a potential causal core gene. To further explore the spatial expression pattern of this gene in the brain, we used PrediXcan^19^ to predict its expression level in each individual of our inhouse sample, to the extent that its expression level in the brain is explained by the genotypes of eQTLs (genetically regulated expression, GReX). The appendix (pp 1) briefed the details of PrediXcan. In our study, we used the pre-stored predictive weights for the expression of *ZNF391* in the 11 brain regions from GTEx (https://gtexportal.org) and the dorsal lateral prefrontal cortex (DLPFC) from the Common Mind Consortium (CMC)^20^, to infer *ZNF391* GReX of participates in our in-house sample. The appendix (pp 2-5) list the recruitment, inclusion and exclusion criteria of participants, quality control and imputation of genomic raw data. All participants signed the consent form, and the study was proved by the ethic committee of West China Hospital, Sichuan University.

Using the linear regression in R, we then identified the brain regions with differentiated expression of *ZNF391* between the cross-disorder patients and the controls (reference level), with the first three population PCs and genotyping batch effect included as covariates.

Following the identification of regions with differential gene expression, we employed a method, t-distributed stochastic neighbour embedding (t-SNE) ^21^, for the clustering of patients according to their spatial pattern of *ZNF391* GReX (biotypes). The methodological details of t-SNE are briefly described in the appendix (pp 5). Besides, we also carried out a K-means clustering analysis to validate the number of clusters arising from t-SNE. We then compared the demographic, cognitive, and brain structural characteristics between different biotypes. Details regarding the cognitive paradigm, MRI scanning, imaging acquisition, preprocessing and statistical methods, are provided in the appendix (pp 5-6).

### Mendelian randomization (MR) analysis

The previous step uncovered significant differences between *ZNF391* biotypes in GMV in RIFOG, and in working memory measures, as assessed by Delay-matching-to-sample (DMS) task of the Cambridge Neuropsychological Test Automated Battery (CANTAB). To further test the causal relationship from gray matter volume (GMV) in right inferior frontal orbital gyrus (RIFOG) to working memory, we adopted an MR approach by setting the *ZNF391* biotype as the instrumental variable (IV) to conduct a path analysis with three multiple regression models as shown below, followed by the estimation of causal effect using a two-stage least square method ^22^.

*ZNF391* biotypes ↬ RIFOG GMV

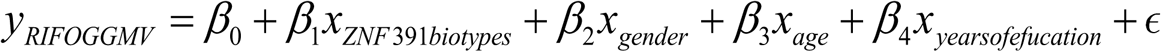

*ZNF391* biotypes ↬ working memory

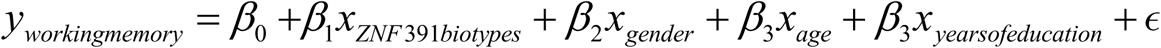

*ZNF391* biotypes ↬ RIFOG GMV ↬ working memory (Empirical)

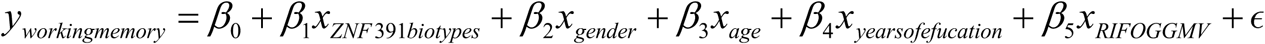

## Results

### Identification of common factor (CF) and ZNF391 as its causal core gene

As illustrated by the pathway diagram (Figure S2; appendix), the genomic SEM of SCZ, BD and MDD revealed a common factor (CF). The loadings of CF on the SCZ, BD and MDD were 0.42, 0.35 and 0.09, respectively, with a comparative fit index (CFI) and a standardized root mean square residual (SRMR) indicative of a well-fitted model (CFI = 1, SRMR = 1.62 × 10^−9^). Besides, consistent with previous studies^23^, the covariance between SCZ and BD is the largest among these three disorders (0.15).

One genomic region in chromosome 6 (262,663,11-293,566,87), containing 39 SNPs, were mapped at a genome-wide significant level (5 × 10^−8^, the Manhattan plot displayed in Figure S3a in the appendix. The top SNP, rs2232429, is located at an intronic region of *ZSCAN12* (P =2.06 × 10^−8^, Figure S4 in the appendix). Besides, no SNP was found to have significant heterogeneity, with the Manhattan plot of Q values for each SNP displayed in Figure 3b in the appendix. Of note, the tissue enrichment analysis using MAGMA in FUMA exhibited a remarkable enrichment of the associated genes in the brain (Figure S5 in the appendix).

**Figure 2:**
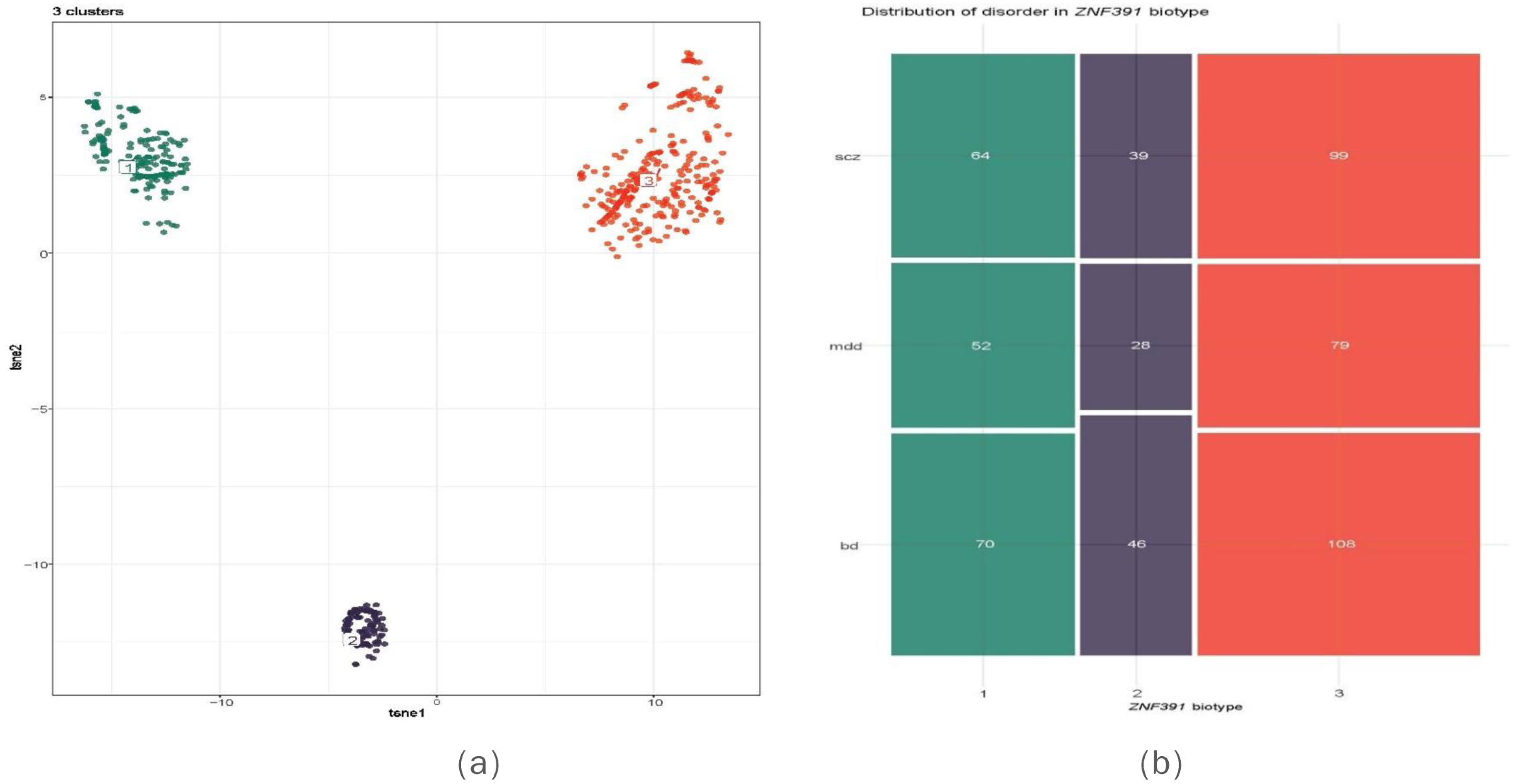
(a) t -SNE results on the transformed two -dimensional dataset with perplexity parameter of 120; (b) the distribution of disorders in each ZNF391 biotype

**Figure 3:**
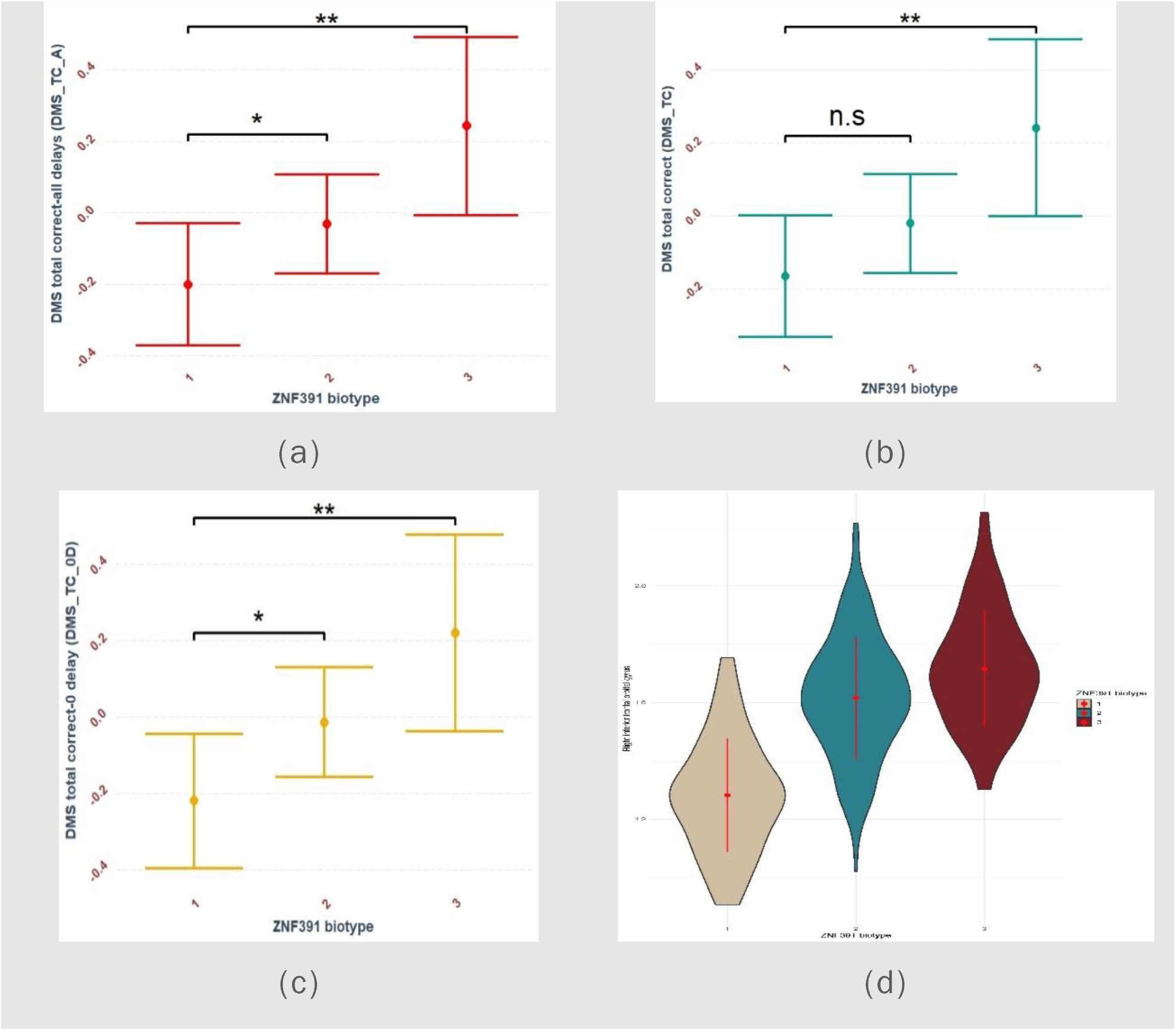
The comparisons of cognitive measures in delay matching to sample (DMS, a-c) and gray matter volume (GMV) of right inferior frontal orbital gurus (RIFOG, d) between three *ZNF391* biotypes

Besides, the SNP heritability of CF was estimated to be 0.2071 (0.0144) and the subsequent partitioned heritability analysis detected the Bonferroni-adjusted (0.05/75 = 6.66 × 10^−4^) significant enrichment of heritability in four cell types (Figure S6; appendix) : Nucleotide_Diversity_10kbL2_0 (P__enrichment_ = 3.39 × 10^−10^), GERP.NSL2_0 (P_ _enrichment_ = 9.51 × 10^−7^), MAF_Adj_LLD_AFRL2_0 (P__enrichment_ = 1.73 × 10^−5^) and Backgrd_Selection_StatL2_0 (P__enrichment_ = *5*.*79* × *10*^−*5*^). The genetic correlation results indicated that the CF was mainly correlated with psychiatric, cognitive and behaviour traits (Figure S7; appendix)

The results arising from *in silico* fine-mapping are summarized in Table S2 in the appendix. The rs7746199-*ZNF391* pair was prioritized with the highest consensus PP (0.96). Based on the calculation and visualization from a centralized platform using S-PrediXcan (https://phenviz.navigome.com/gene_phenotypes/ENSG00000124613.html) ^24^, the inferred expression of *ZNF391* decreases by at least two standard deviations in the mental and behavioural traits (mainly schizophrenia, Figure S8, Figure S9 in the appendix).

### Brain regions indicative of a differentiated ZNF391 expression and novel biotypes arising from differentiated expression

After the sample quality control and population structure analysis, we included 585 patients (202 SCZ, 224 BD, 159 MDD) and 386 healthy controls, each with 6,230,803 high-quality SNPs which passed genotype quality control and remained in the subsequent analysis for the inference of *ZNF391* GReX (demographic characteristics in Table 1, and consort diagram for the quality control of genotypes in Figure S1 in the appendix). As displayed in Figure S10 in the appendix, no population stratification was detected in our sample, and all 971 individuals were of Han Chinese ancestry, clustering in the East Asian population. As shown in Figure S11 and Table S3 in the appendix, 8 out of 12 tested brain regions showed a significant difference in *ZNF391* GReX between patients and controls, with the most significant difference observed in the frontal cortex (P = 0.003). It is worth mentioning that we did not find the *ZNF391* GReX significantly different between disorders.

**Table 1:**
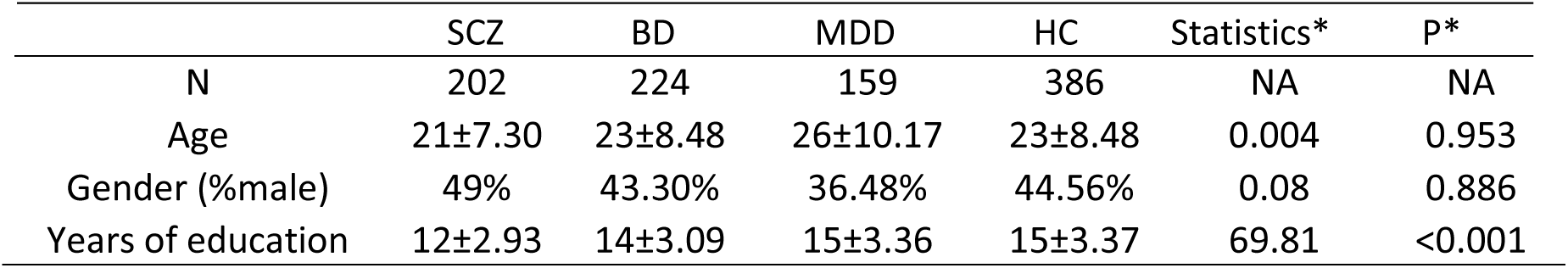
Demographic characteristics of the participants involved in the inference of *ZNF391* GReX; *denotes the statistics and P-values derived from the comparison between the patients from three disorder groups (SCZ + BD + MDD) and controls

The clustering analysis based on the spatial expression pattern of *ZNF391* GReX indicated that three biotypes (Figure 2a) could be detected in our patient group in the most optimal condition (perplexity = 120, Figure S12 in the appendix). A χ^2^ test did not find a significant difference in the distribution of disorders between three biotypes (Figure 2b). The results using K-means clustering analysis overlapped 100% with the result from t-SNE.

No significant difference was found in age, gender and years of education between three biotypes of *ZNF391* GReX. Comparison of cognitive function using CANTAB showed that working memory in the paradigm of Delay-matching-to-sample (DMS) was the only domain to exhibit differential performance between biotypes; as shown in Figure3(a-c), after regressing out the effect of age, gender and years of education, biotype 3 group performed significantly better than other 2 biotype groups (biotype 3 > biotype 2 > biotype 1) in the DMS total correct (DMS_TC), the DMS Total correct-all delays (DMS_TC_A), and the DMS total correct-0 delay (DMS_TC0D). The comparison of GMV between different biotypes identified the GMV in RIFOG to be significantly different between three biotypes, mirroring the order of DMS comparison results (P_anova_ = 0.0027, Figure 3d).

### *A partial causal relationship existing between* GMV of RIFOG and working memory

With *ZNF391* biotypes as an IV, a causal path, albeit a partial one, could be drawn from RIFOG GMV to working memory **(β**_DMS_TC0D_ = 4.95, P = 0.0056; **β**_DMS_TC_ = 2.53, P = 0.059; **β**_DMS_TC_A_ = 2.57, P = 0.056). As demonstrated in Figure 4, the signal of association with *ZNF391* GReX decreased gradually from gray matter volume to all three measures of working memory; while the hypothetical total effect of pathway (*ZNF391* GReX ↬ RIFOG ↬ working memory) was estimated to be 2.24% on average, the empirical estimation based on a linear model implied an effect around 8.3%, emphasizing the complexity of the biological mechanism linking gene to the complex trait (Figure 4).

**Figure 4:**
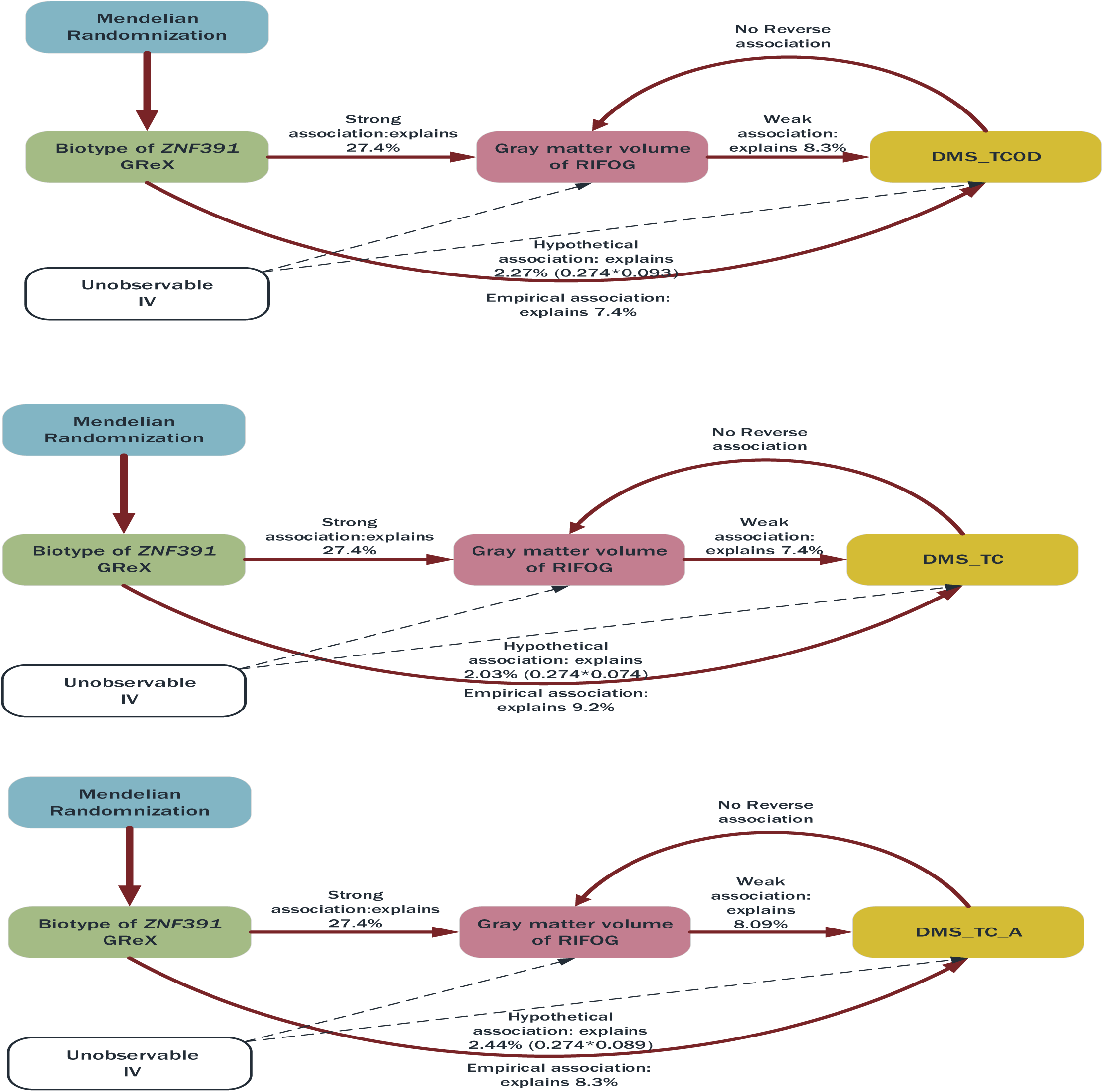
Application of Mendelian randomization for identifying the causal relationship between GMV of RIFOG and working memory

## Discussion

Contrary to the conventional bottom-up study design, we took a top-down approach by distilling signals from the large-scale GWAS of psychiatric disorders for further exploration in our sample. We argue that such an approach could both reduce the risk of false-positive discovery (“Winner’s curse”) and provide the robustness for the investigation of the biological mechanism at the higher level of granularity. Compared to a polygenic risk score (PRS), PredixCan is affected by the population allelic difference in a more attenuated way in that, instead of using OLS as the training algorithm, PredixCan chose a sparse algorithm, elastic net, which, could improve predictive efficacy. Further, Mogil et al. ^25^ showed that best population-specific predicted gene has a highly correlated performance across populations, echoing the findings from another study that 83% of genes differentially expressed among individuals, and 17% differentially expressed among populations, with the most variation coming from within-population ^26^. All these studies justify the application of expression-level data in the across-population studies.

Consistent with previous studies ^5^, our study detected a common factor (CF) underlying three psychiatric disorders (SCZ, BD and MDD), and the CF was associated with a segment at the extended major histocompatibility complex (MHC) region. Sekar *et al.* identified the variations of C4 in the MHC region as the driving signal of association with schizophrenia; when looking beyond the MHC region, Sekar *et al.* detected a pattern of bimodal associations in the sense that two peaks of association existed in this region, one was C4, another was the extended region harbouring the signal of interest in our study, *ZNF391* ^27^. *ZNF391* is highly expressed in the brain, was implicated in depressive phenotypes ^28^, schizophrenia ^29^, neuroticism ^30^. To the best of our knowledge, this is the first study to dissect the role of *ZNF391* at a different level of granularity, from expression to gray matter volume (GMV) and working memory. Carrying out a genetic expression inference analysis by using our in-house data, we detected eight brain regions with a differentiated expression of *ZNF391*. Of these eight brain regions, six are subcortical regions, two are cortical regions, which echoed widely recognized knowledge that psychiatric disorders implicated many brain regions^31^.

Instead of focusing on comparing patients with healthy controls, our study took the results to the next level by defining new biotypes in the patient group based on the expression pattern of *ZNF391* in these eight brain regions. To date, we believe our study is the first one to define the new biotypes of psychiatric disorders by incorporating the *in silico* genetic expression data and clustering algorithm. Although future replication using an independent sample is needed, our study provides an alternative approach to evaluate the *in vivo* gene expression in the tissues implausible to collect, such as the brain. Besides, with *ZNF391* GReX as IV, we demonstrated a partial causal path from GMV of RIFOG to working memory, which vividly illustrates widespread horizontal pleiotropy in the human genome ^32^. Genetic instruments, from which the term “Mendelian Randomization” was coined, gained popularity gradually as IVs due to their determination by Mendelian segregation, being non-modifiable (except for somatic mutations) ensuring lifelong exposure and mitigating concerns about reverse causation^33^. While the genetic IVs could aid in making complete causal inference in the condition underpinned by one or a few genes, they provided limited information in the complex traits due to the small effect size of each of these IVs. In our study, we chose GReX as IV, which not only could make a causal inference, but also provide insights into the proportions of phenotypic relationship attributable to specific genes and pathways ^34^.

### Strengths and Limitations

The greatest strength of present study lies in the fact that we used the signal from the large-sample GWAS. Such a strategy ensures the robustness of the signals we used for our subsequent analysis, and our results, in return, could enrich our understanding of these signals in a more refined way. Another strength of our study is that we used a tissue-specific approach to explore further the signal, which narrows the gap between gene and its phenotypic consequences.

The current study should be interpreted in light of following two limitations: (1) we used the largest available gene expression project, GTEx, in our study to infer GReX and we are fully aware that most of the donors in GTEx are of European ancestry. Although the power lost to the allelic frequency difference due to population structure is more attenuated than that in PRS calculation, we are fully aware of the lesser degree of the power loss. This discrepancy in predictive accuracy underscores the importance of adding samples from diverse populations to the current database for the success of understanding the biology behind genetic variations, ultimately leading to the success of precision psychiatry. (2) Meanwhile, there might be a sampling bias in our dataset, the patients recruited in our study who were able to finish the evaluation including cognition test and MRI scanning might not represent the whole population of patients.

### Conclusion

Taken together, in our current study, we tried to define new cross-disorder biotypes by using a cross-disorder top-down approach. The findings of this study suggest a common pathological mechanism, to a different extent for each disorder, may underlie the three disorders. Our results showed that a general liability underlies SCZ, BD and MDD, with *ZNF391* being a potential causal core gene conferring risk of such a general liability. Moreover, we re-clustered patient group into three biotypes based on their expression profiles of *ZNF391* in the brain. The subsequent analysis led to the linkage between *ZNF391*, working memory and gray matter volume (GMV) of RIFOG. Although future studies are required to delve deep into the biological mechanism linking them together, our study provides an example vis-à-vis how to increase the granularity of genetic study to further our understanding of etiology of psychiatric disorders by incorporating the knowledge from the big data and prediction algorithm into the real-world data.

## Supporting information

Supplementary online content

## Funding

The National Key Research and Development Program of the Ministry of Science and Technology of China; National Nature Science Foundation of China Key Project; 1.3.5 Project for discipline of excellence, West China Hospital; Science and Technology Program of Guangdong

### Contributors

HYR designed, analyzed and drafted the manuscript; YJM & YMZ processed neuroimaging and neurocognitive data; QW and WD co-supervised the study; XJL,LSZ and YCW collected the data; XHM,PS and TL are principal investigators and supervised the study.

