## Supplementary online content for "Spatial expression pattern of *ZNF391* gene in the brains of patients with schizophrenia, bipolar disorders or major depressive disorder identifies new cross-disorder biotypes: A trans-diagnostic, top-down approach"

**Appendix**

1. **Methodological details of genomic SEM**

Genomic SEM takes the genetic covariance matrix estimated from linkage disequilibrium (LD) score regression algorithm and fits a structural equation model to it; we specified an SEM containing a common factor (CF) which, to different extents, simultaneously explain the phenotypic variances and covariances of the three disorders. Compared to other multivariate GWAS methods, Genomic SEM, instead of arbitrarily deriving a multivariate effect for each SNP, is able to model shared genetic architecture across phenotypes with customized factors and compared different models ^1^.

2. **Methodological details of *FINEMAP and iRIGS***

The first algorithm, “FINEMAP” ^2^, is a stochastic search algorithm for identifying the casual configurations of SNPs with substantial posterior probability (PP) in the genomic region of interest. Considering that the significant association could arise from between one eQTL and the multiple genes in the multiple tissues, we chose the gene regulated by the causal SNP and also highly expressed in the brain according to the GTEx data repository, as the target gene. The second, iRIGs ^3^, is a Bayesian framework which infers PP for the causal SNP-gene pair by integrating two layers of information: (1) evidence from multi-omics data, and (2) relationships among genes in biological networks. In our study, we define the SNP-gene pair that reached a consensus PP of 0.9, obtained by averaging PPs from the two algorithms, as being causal.

3. **Methodological details of PrediXcan**

In brief, the PrediXcan first used the gene expression data in different tissues and the genotype data from the same dataset to train the elastic net models, with the expression level of the gene as the response variable and the genotypes of each SNP as the primary predictor variable. These models generated, for each SNP, a regularised weight of prediction. Subsequently, using genotype data from an independent sample, the predictive weights of SNPs within 1MB of the gene start or end (according to the GENCODE version 12) were aggregated using the following formula (1) to yield GReX for the target gene in the target tissue:

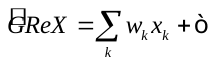
 (1)

In the equation above, X_k_ is the number of reference alleles for SNP k.

**4. Recruitment, inclusion and exclusion criteria of involved participants**

The patients with schizophrenia were first-episode, drug-naïve at the time of evaluation; the patients with major depressive disorder were either first-episode, drug-naïve or had stopped taking antidepressant, mood stabilizer or other psychotropic medications for at least three months prior to the evaluation; we included the patients with bipolar disorder regardless of their medication status at the time of assessment due to availability of sampling. The inclusion and exclusion criteria are listed below:

*For first-episode schizophrenia:*

(1)	Aged between 16 and 55

(2)	Han Chinese

(3)	Right-handed

(4)	Experience the first episode of psychosis at the time of recruitment

(5)	Fulfill the diagnosis criteria of schizophrenia in the Diagnostic and Statistical Manual of Mental Disorders (DSM-IV)

(6)	IQ >= 70 according to Wechsler IQ Test

(7)	Voluntarily participate in the current study with a written consent form at the time of evaluation

(8)	The current episode cannot be accounted for by any specific life events.

*For bipolar disorder, the inclusion criteria are the same as that of schizophrenia except for the item (1), (4) and (5):*

(1) Age between 12 and 55,

(4) Either at the first episode or the recurrent episode of bipolar disorder

(5) Fulfill the diagnosis criteria of bipolar disorder in the Diagnostic and Statistical Manual of Mental Disorders (DSM-IV)

*For major depressive disorder, the inclusion criteria are the same as that of schizophrenia except for the item (1), (4) and (5):*

(1)	Age between 16 and 60

(4)	Either at the first episode or the recurrent episode of major depressive disorder, with active symptoms.

(5)	Fulfill the diagnosis criteria of major depressive disorder in the Diagnostic and Statistical Manual of Mental Disorders (DSM-IV)

The patients were interviewed and assessed by trained psychiatrists using the Structured Clinical Interview for the DSM-IV (Diagnostic and Statistical Manual of Mental Disorders, fourth edition) (SCID-I/P). Patients with schizophrenia was followed-up for at least six months for the verification of their diagnoses. Patients would be excluded if they met any of the exclusion criteria below:

(1)	Comorbid with serious organic brain disorder, or neurological diseases or severe endocrinological and metabolic diseases

(2)	Comorbid with other conditions defined with the axis I and axis II disorders, such as substance abuse, intellectual disability

(3)	Have metallic implants at the time of MRI scanning, for example, dentures, pacemaker and prostheses

(4)	Color-blinded or stuttering

(5)	Pregnant or postpartum at the time of recruitment

(6)	Received ECT 6 month before recruitment

For the healthy controls, the inclusion criteria are:

(1)	No indication of any axis I and axis II disorders based on Structured Clinical Interview for DSM-IV, SCID-NP.

(2)	Age between 12 and 60

(3)	Level of education no lower than graduation from elementary school

(4)	Han Chinese

(5)	Right-handed

(6)	IQ >= 70 according to Wechsler IQ Test

(7)	Intact social functioning with full capacity for one’s behaviour

(8)	Voluntarily signed the consent form

The exclusion criteria are:

(1)	History of axis I and axis II disorders based on DMS-IV

(2)	History of being prescribed with antipsychotics, antidepressants and mood stabilizer, the apparent use of benzodiazepine one month before MRI scanning

(3)	Have metallic implants at the time of MRI scanning, for example, dentures, pacemaker and prostheses

(4)	Color-blinded or stuttering

(5)	Pregnant or postpartum at the time of recruitment

5. Genotyping and quality control

In our study, DNA samples of 988 individuals (431 males, 551 females, 6 with missing gender status) were collected and underwent the whole-genome genotyping using Global Screening Array (GSA-24 V2.0) and Infinium OmniZhongHua-8 chips. Among these individuals, 399 are healthy controls, 589 are patients with one of the three psychiatric disorders (schizophrenia: 203, bipolar disorder: 226: major depressive disorder: 160). The quality control of genomic data followed the routine procedures, of note, sex of 6 individuals with missing gender information was imputed using genomic data : SNPs were filtered based on unmatched sex information between genomic sex status and self-reported gender information, missing genotype rate (>= 3%), heterozygosity (>= 3rd standard deviations), Hardy-Weinberg equilibrium (P < 0.001) and common frequency (MAF > 0.01); the filtering of individuals is illustrated in the Figure S1. Following the systematic quality controlling, we chose haplotype reference panel (HRC) on the platform of Michigan Imputation Server to impute missing genotypes, followed by a brief quality control steps including the filtering of imputation quality (info >= 0.3), missing genotype rate (>= 3%) and major allele frequency (MAF > 0.01). In total, 971 individuals, each with 6,230,803 high-quality SNPs, passed the quality control and remained in the subsequent analysis. As displayed in the Supplementary Fig2(a) and 2(b), no evident population stratification exists in our sample, and all 971 individuals are of Han Chinese ancestry, clustering in the East Asian population. Quality control was implemented using PLINK 1.9 ([www.cog-genomics.org/plink/1.9/](http://www.cog-genomics.org/plink/1.9/)) ^4^, and population stratification analysis was carried out using EIGENSTRAT ^5^.

6. Clustering analysis of ZNF391 GReX using t-SNE

T-SNE uses random walks on neighbourhood graphs to allow the implicit structure of all data to influence how a subset of the data is displayed. Compared to many other similar non-parametric techniques, t-SNE can better capture much of the local structure of the high-dimensional data while also revealing global structure such as the presence of clusters at several scales^6^. Before the clustering, the confounding effect of population stratification and the genotyping batch were regressed out. We iterated on different values of its super parameter, perplexity (range 50-150) to find the most optimal value based on the within-cluster closeness inspection, and also carried out a K-means clustering analysis to validate the primary t-SNE clustering findings.

7. Cognitive comparison using CANTAB, imaging acquisition and preprocessing

we chose the paradigms from the Cambridge Neuropsychological Test Automated Battery (CANTAB) with its primary outcomes and cognitive domains of CANTAB listed in the Table S1.

High-resolution T1-weighted images were acquired from subjects on admission using a 3-Tesla MRI system (Achieva; Philips, Amsterdam, the Netherlands) and an eight-channel phase array head coil with a volumetric 3D Spoiled Gradient Recall (SPGR) sequence (TR=8.1ms, TE=3.7ms, slice thickness=1.0mm [no slice gap], FOV=24×24 cm2, matrix=256×256), generating 188 contiguous axial slices with the in-plane resolution of 1mm×1mm. During scanning, all participants were ears plugged, foam padded, instructed to remain motionless through the whole scanning process. Two experienced radiologists inspected the quality of the raw image data. No gross graphic abnormalities were detected in any participant.

Structural data were preprocessing using voxel-based morphometry (CAT12: Computational Anatomy Toolbox 12) and SPM12 (http://www.fl.ion.ucl.ac.uk/spm) running on Matlab. Then, we had acquired the gray matter volumes (GMV) at the different brain regions according to the Neuromorphometrics Atlas implemented in SPM12. For all VBM analyses, we included total intracranial volume (TIV) and GMV as a covariate to remove variance related to this global parameter of brain morphometry. We then performed whole-brain voxel-wise analyses calculating both positive and negative correlations between regional brain volumes and each of the (sub) scales separately.

8. Statistical analysis

A Kruskal-Wallis test was used to compare the mean of continuous demographic features, including age and years of education, between three biotypes; categorical features, including gender and disorder distribution, were tested using x2 test. Univariate analysis of variance was conducted for the comparison of cognitive performance and gray matter volumes (GMV), both cognitive test and GMV were adjusted for age, gender and years of educations before comparison. All of the statistical analysis was carried out in the R, version 3.5.3 (R foundation). Given the high correlation between measures of domains in CANTAB, we chose a liberal threshold of α =0.05 as the significance level.

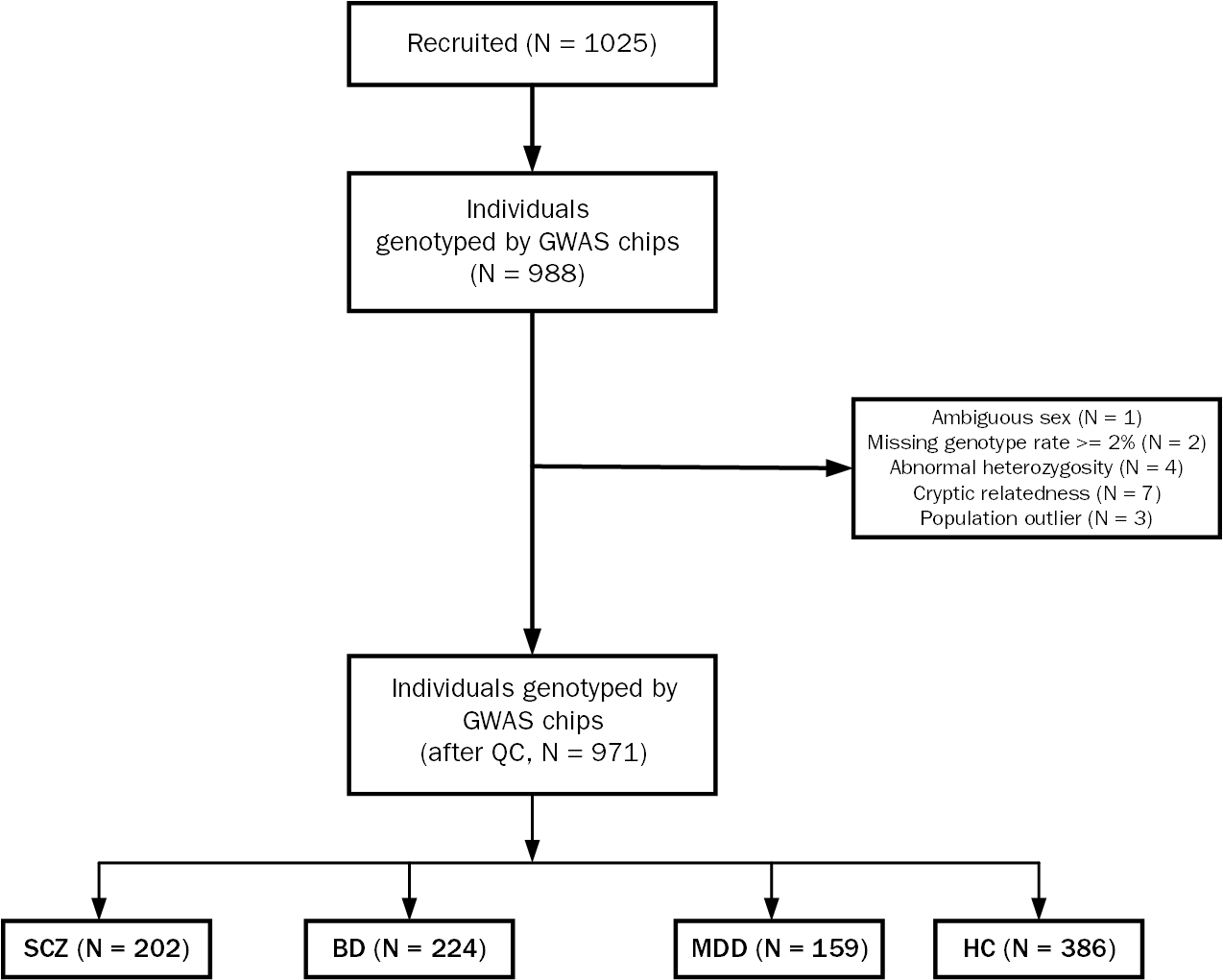

Figure S1. Quality control steps to filter genotyped individuals

| Dimension of cognitive function | Test | Abbreviation | Main outcomes |
| --- | --- | --- | --- |
| Visual working memory test | Delayed matching to sample | DMS | 1.Latency 2.Numbers and percentages correct, and errors 3.Signal detection theory measures |
|  | Pattern Recognition Memory | PRM | 1.Numbers and percentages  correct and  incorrect 2.Latency |
| Executive function and planning test | Intra-Extra dimensional SetShift | IED | 1.Errors 2. Numbers of trials and stages (blocks) completed |
|  | Spatial working memory | SWM | 1.Errors (between, within, double, and total)  2. Strategy  3. Latency  4. Problem reached |
|  | Stockings of Cambridge | SOC | 1.Problems solved in minimum moves  2. Mean moves  3. Initial thinking time  4.Subsequent thinking time |
| Attention test | Rapid Visual Information Processing | RVP | 1. Hits, misses, false alarms and rejections  2. Probabilities and sensitivity calculated using SDT  3. Latency |

Table S1. Paradigms in CANTAB we used for the cognitive evaluation in the study and their main outcomes

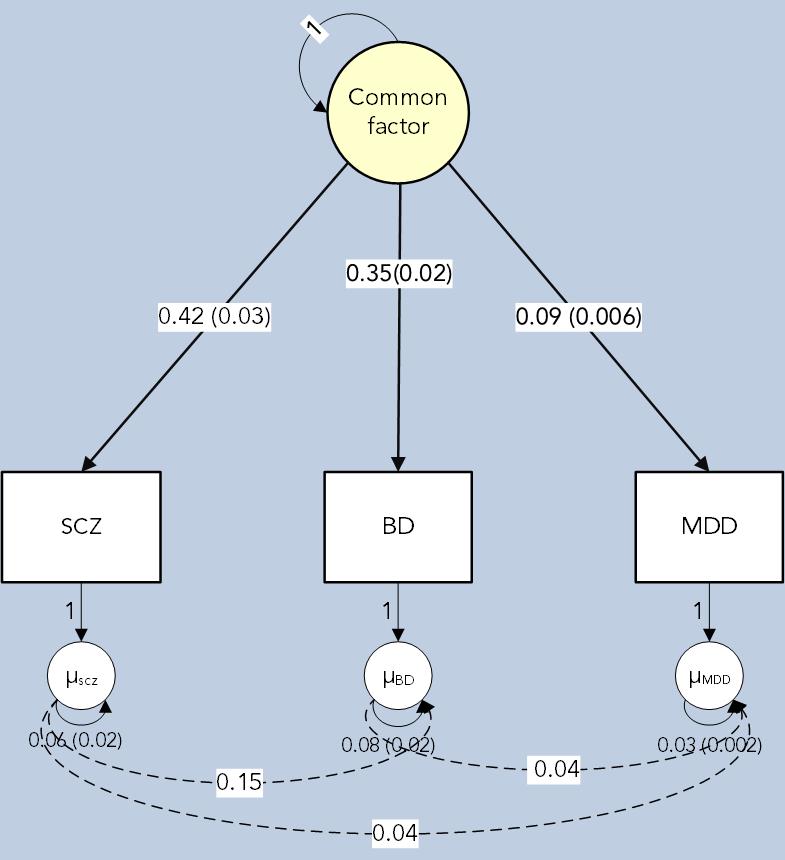

Figure S2. Pathway diagram of the genomic SEM solution for a common factor (CF) underlying schizophrenia (SCZ), bipolar disorder (BD) and major depressive disorder (MDD)

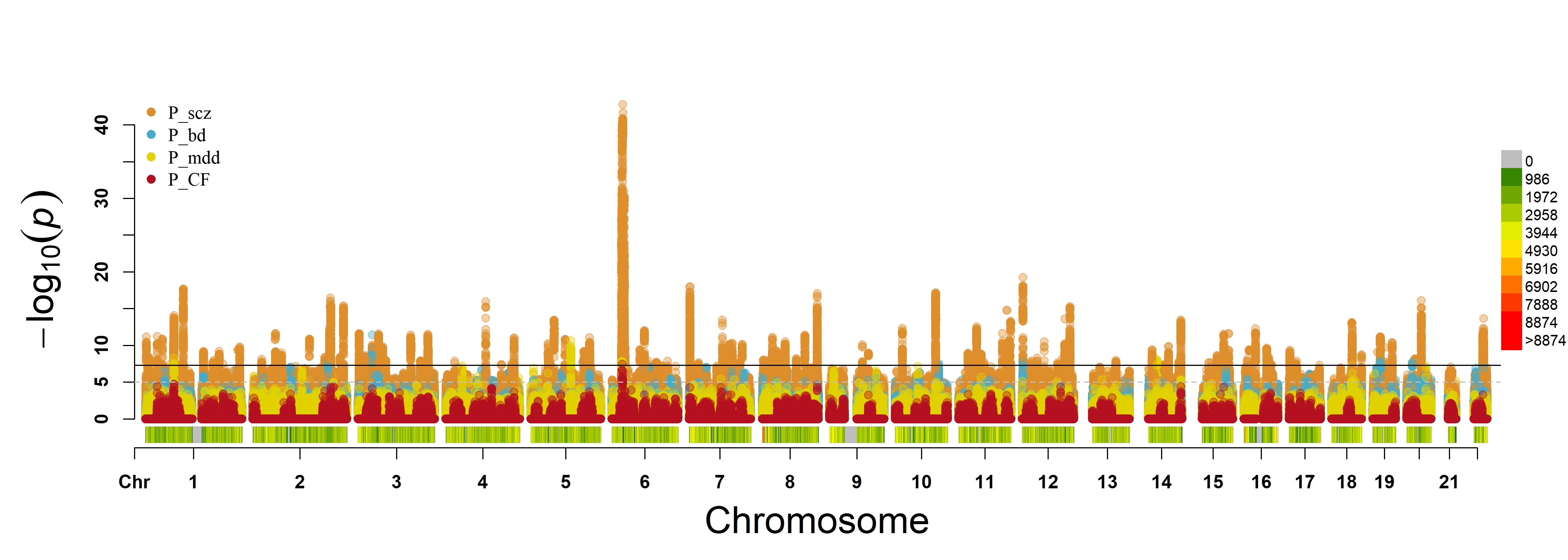

(a)

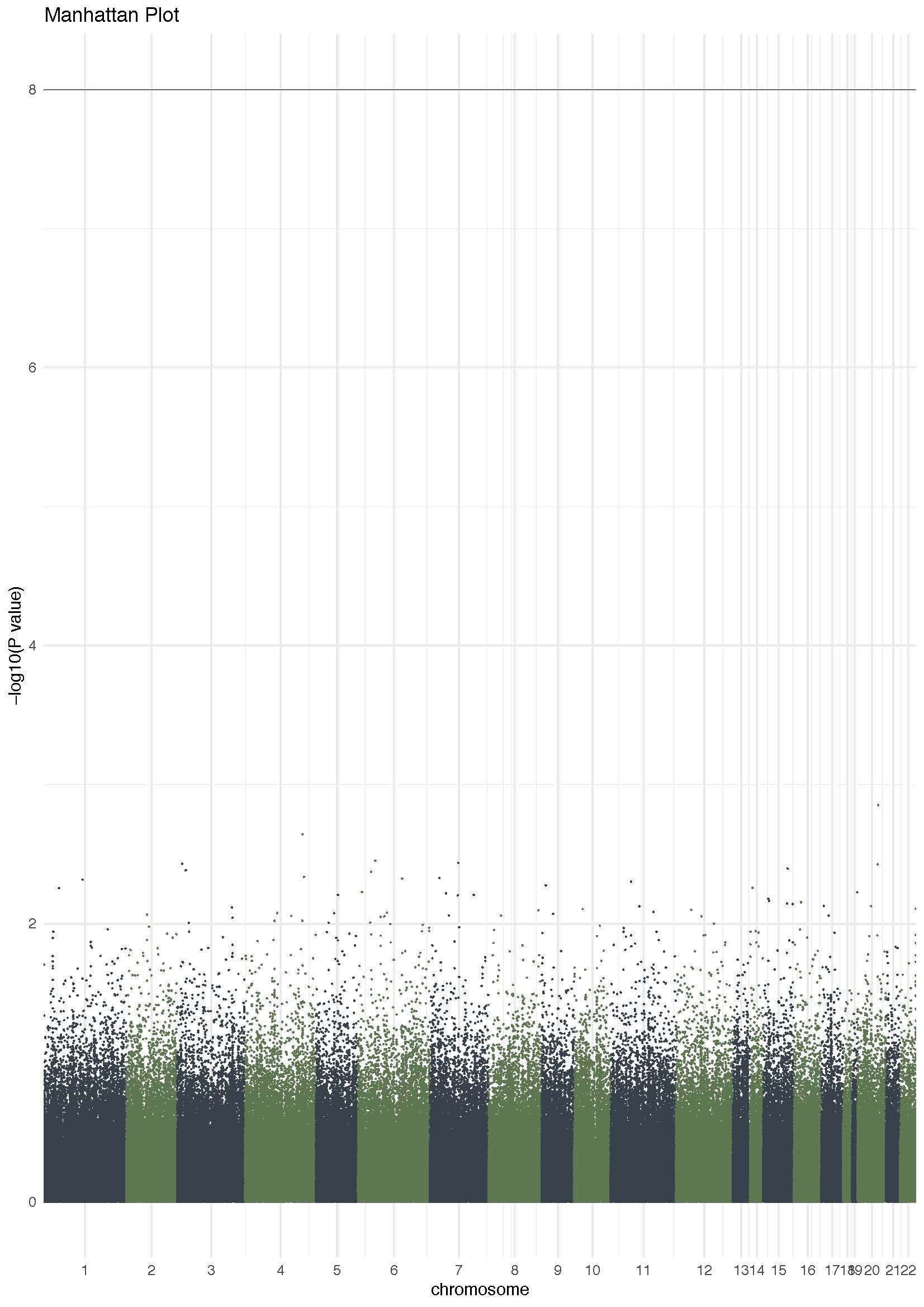

(b)

Figure S3. (a) The Manhattan plot for the composite results from SCZ, BD, MDD and CF GWAS, the ruby points represent the results of CF; the marigold points represent the results of SCZ GWAS by Pardiñas, A.F. et al; arctic points represent the results of BD GWAS by Stahl, E.A et al.; canary points represent the results of MDD GWAS by Wray, N.R et al. (b) Manhattan plot of Q value for each SNP. In both plots, each point denotes the -log10 P-value for each included SNPs, the black line marks the threshold for genome-wide significance (5×10^-8^)

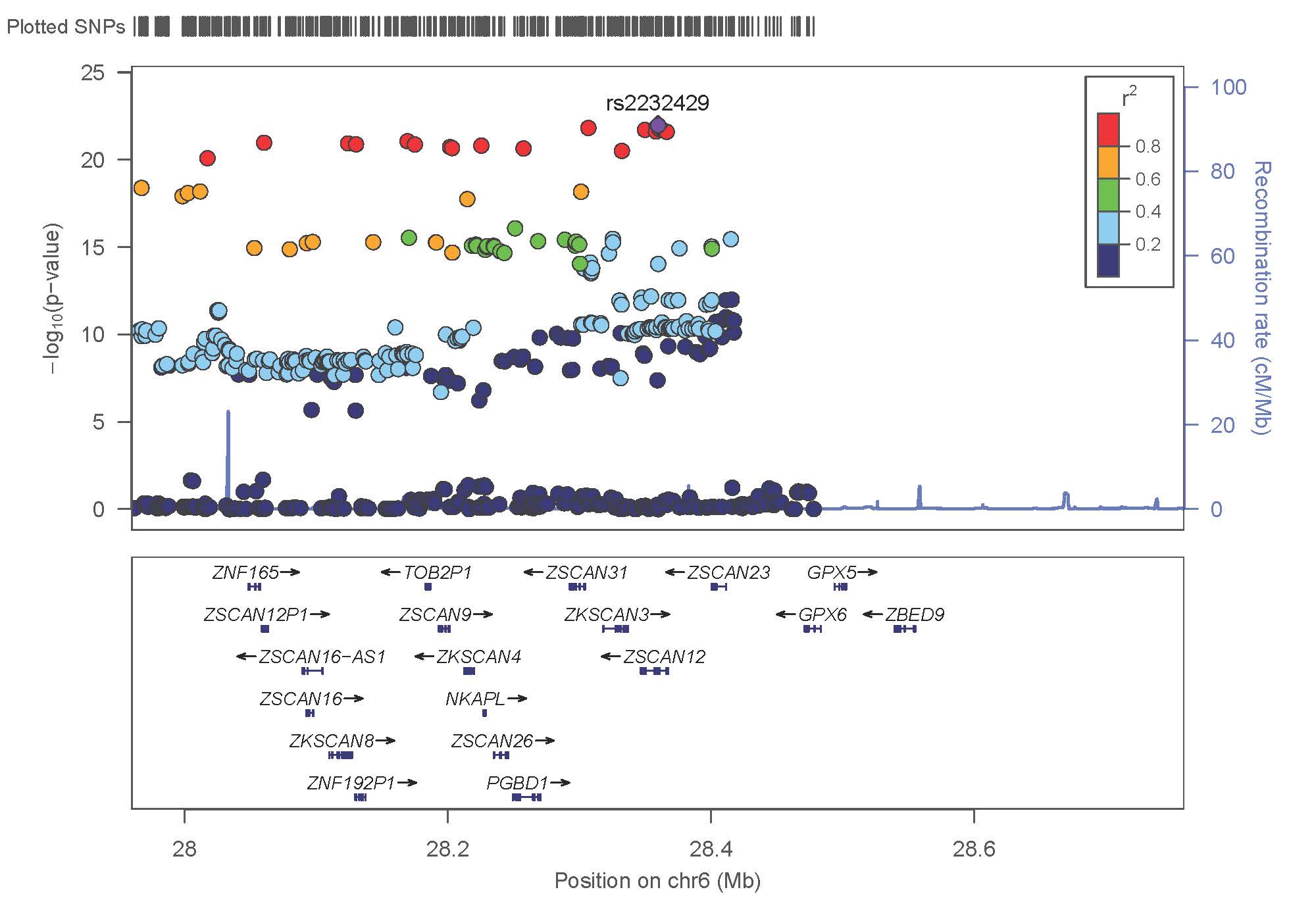

Figure S4. A regional association plot of and its linkage disequilibrium with neighbouring SNPs, the purple diamond denotes rs2232429 which showed the strongest signal of association in CF GWAS

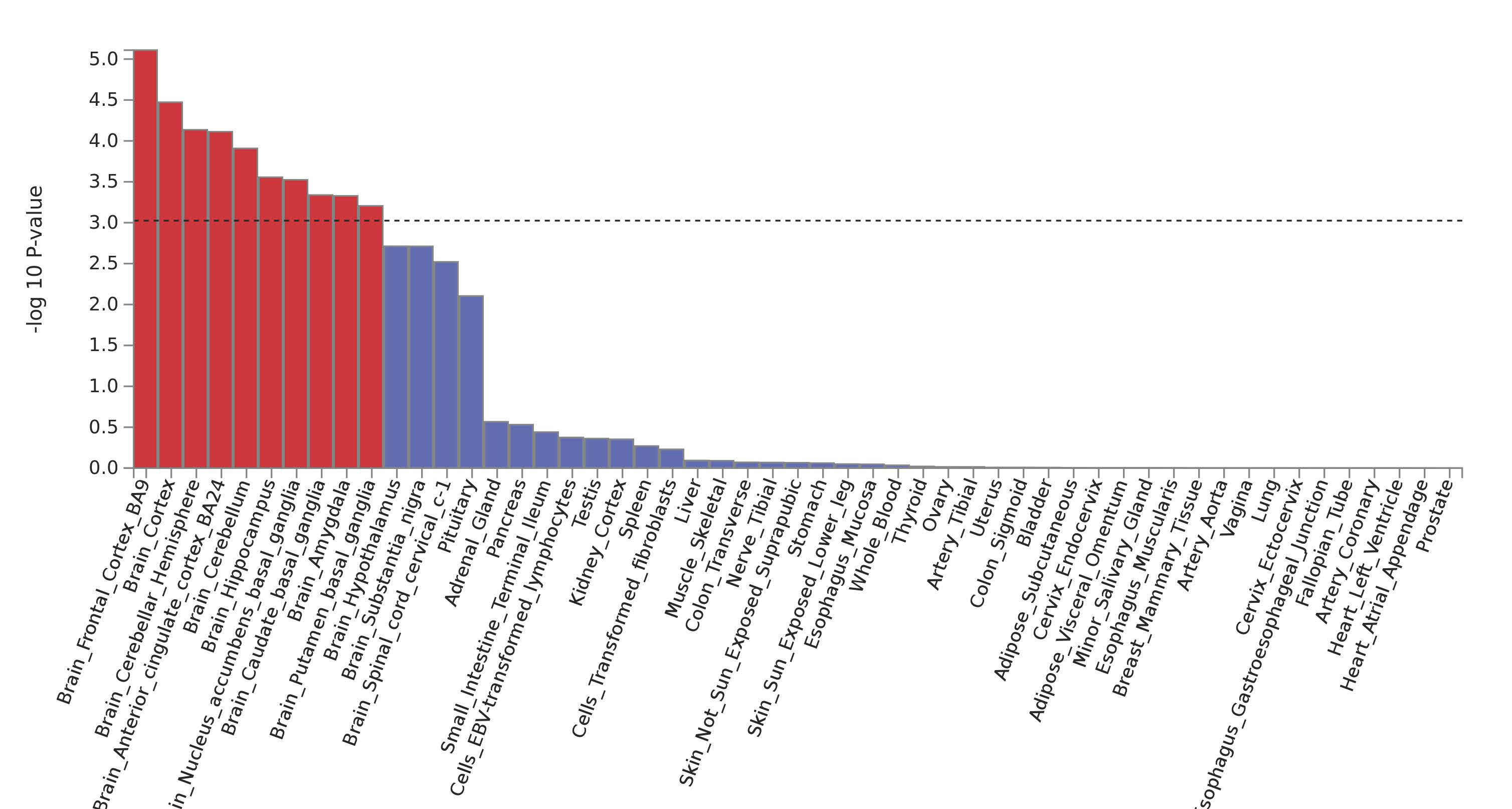

Figure S5. The results of tissue expression enrichment using MAGMA in FUMA; the red bars are the tissues significantly enriched by the genes associated CF and the black dashed line indicates the Bonferroni-corrected significant threshold

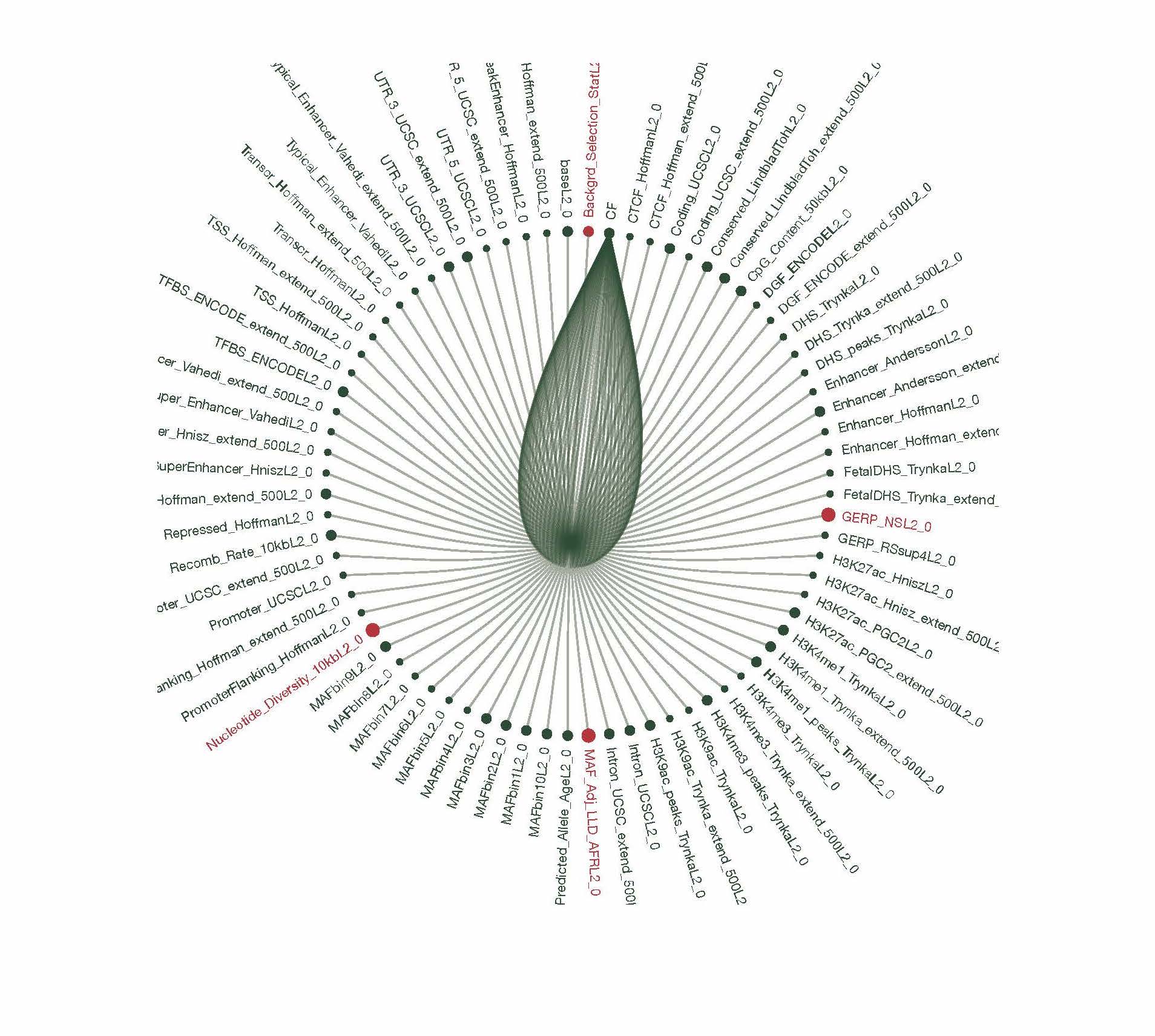

Figure S6. The analysis result of heritability enrichment in different cell types; the red dots represent the cell type contributing significantly to the heritability of CF, with the significance surviving a Bonferroni correction for multiple comparisons

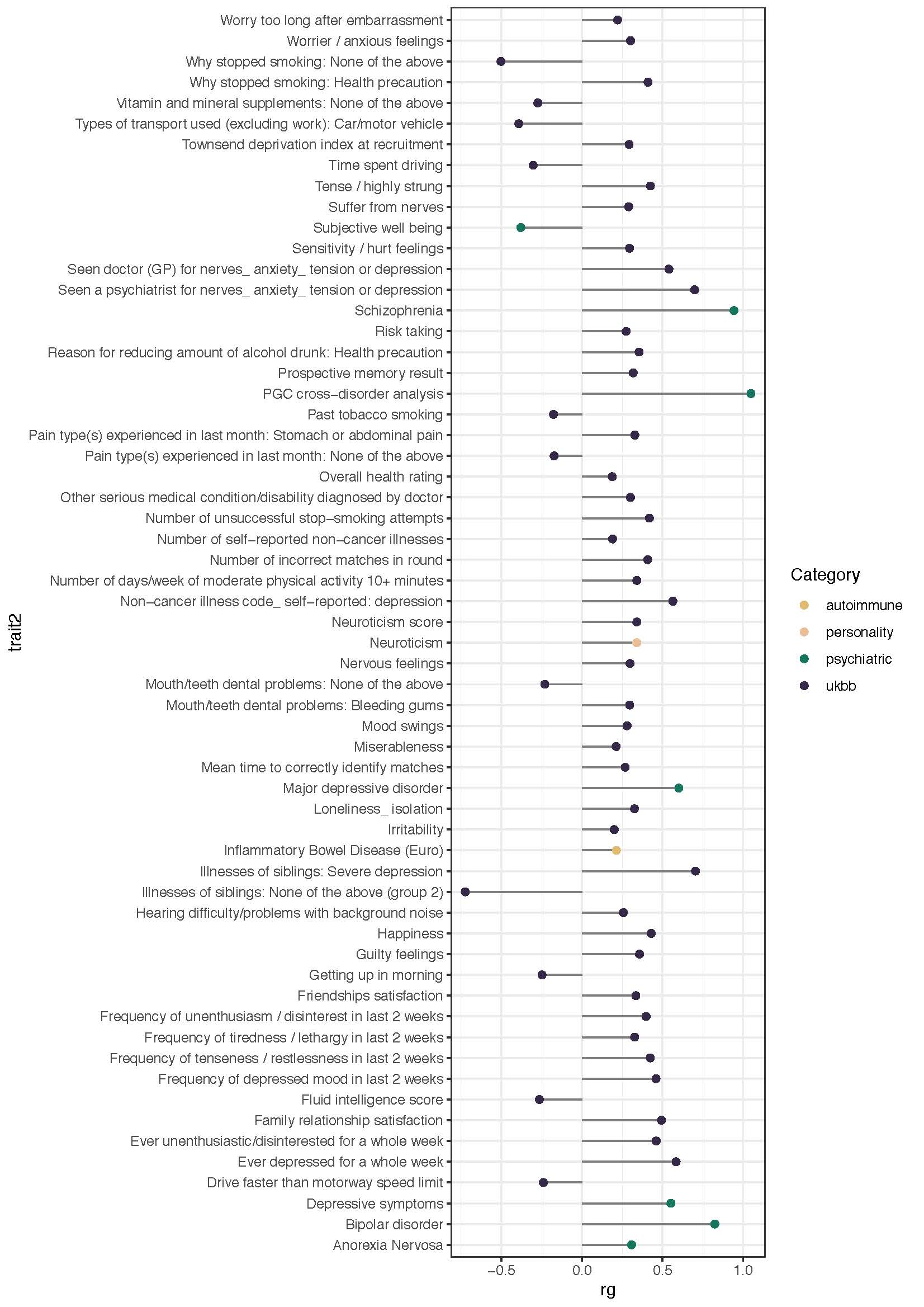

Figure S7. The genetic correlation between common factor and the relevant traits in LD hub database

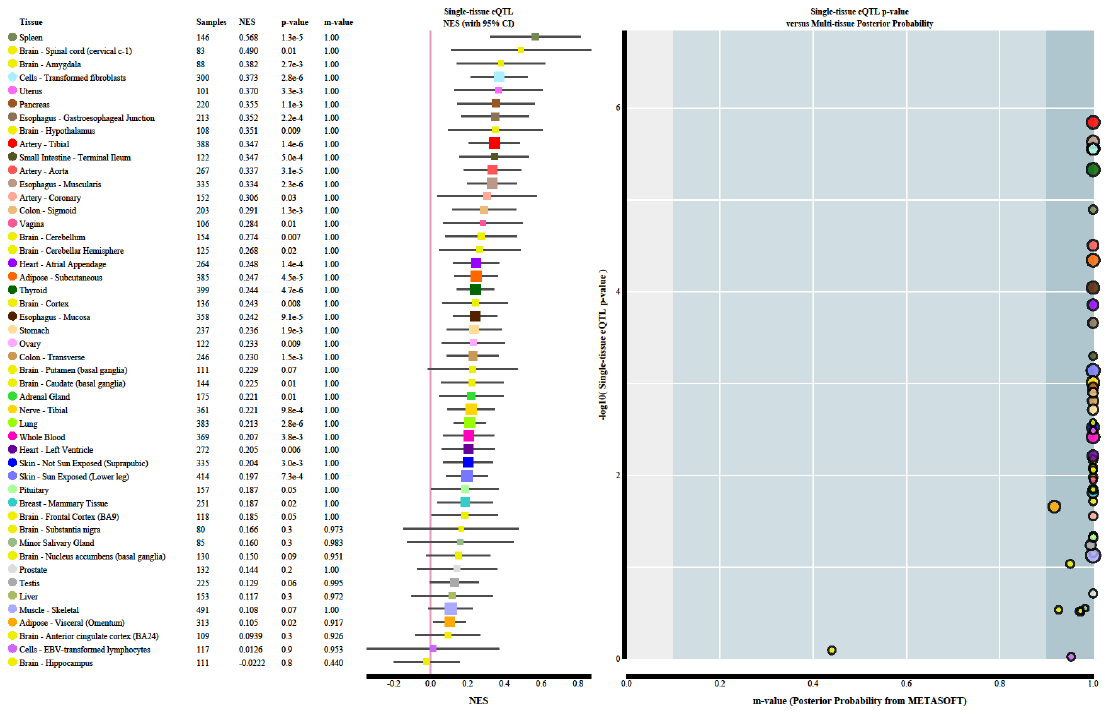

Figure S8. Multi-tissue plot for effect of rs7746199 on ZNF391 and corresponding posterior possibility, downloaded from <https://gtexportal.org/home/snp/rs7746199>

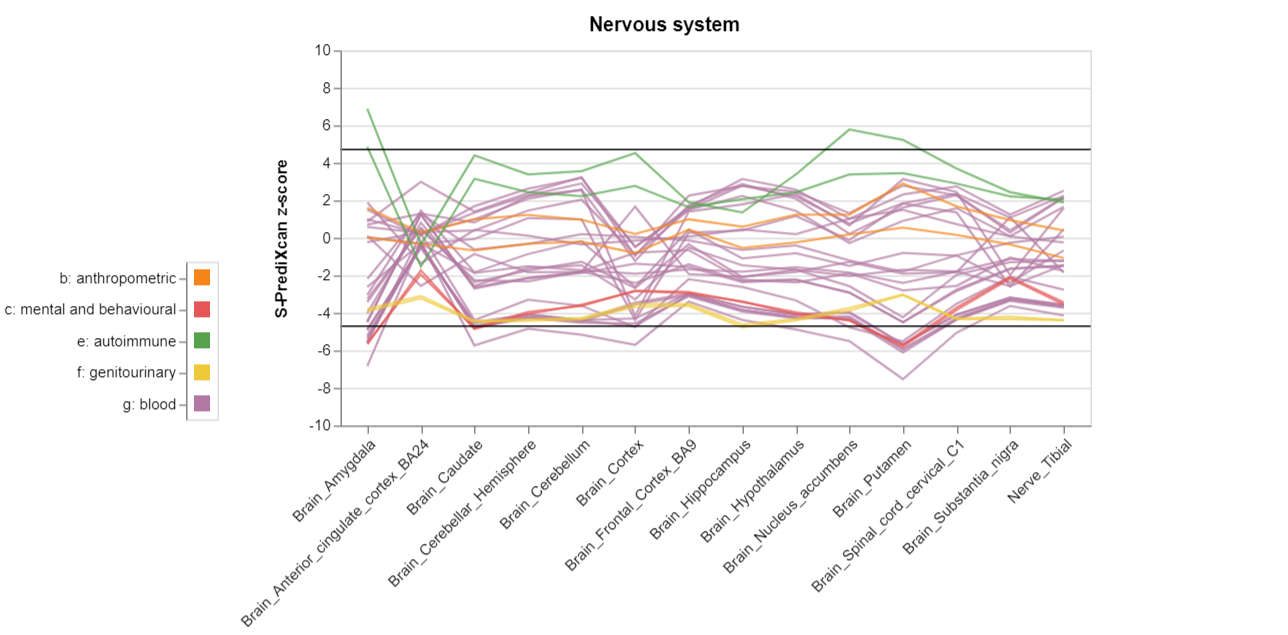

Figure S9: Inferred ZNF391 GReX in the brain regions using the summary statistics of traits of different categories, downloaded from <https://phenviz.navigome.com/gene_phenotypes/ENSG00000124613.html>

| 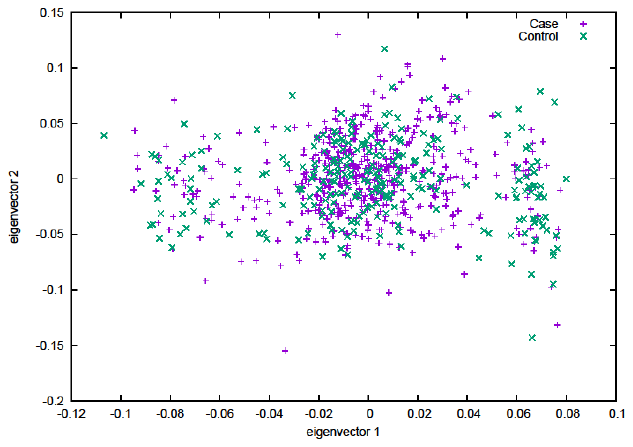 | 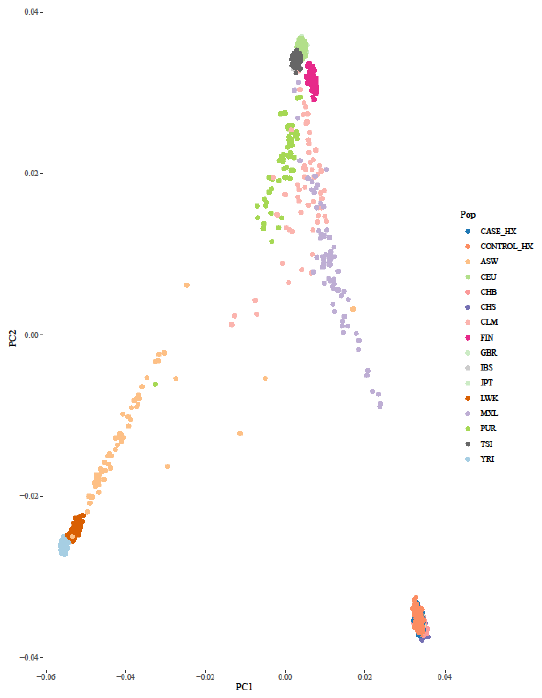 |
| --- | --- |
| (a) | (b) |

Figure S10. Population scatter plot of PC1 and PC2, stratified by disease status (a), and in the context of populations in the 1000 Genomes (b)

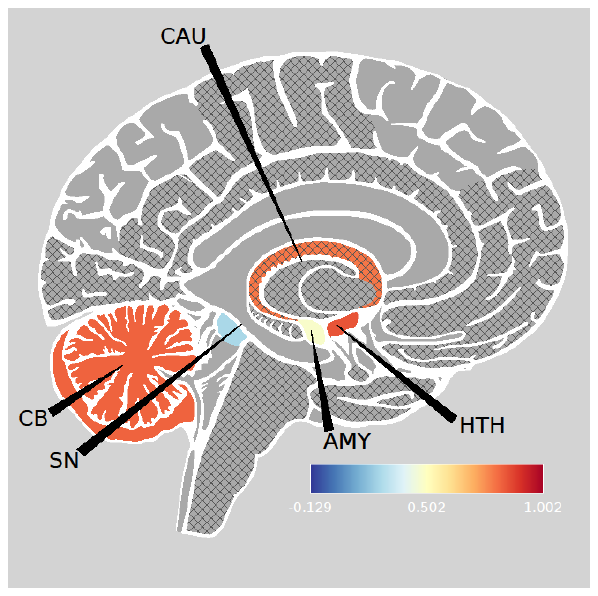

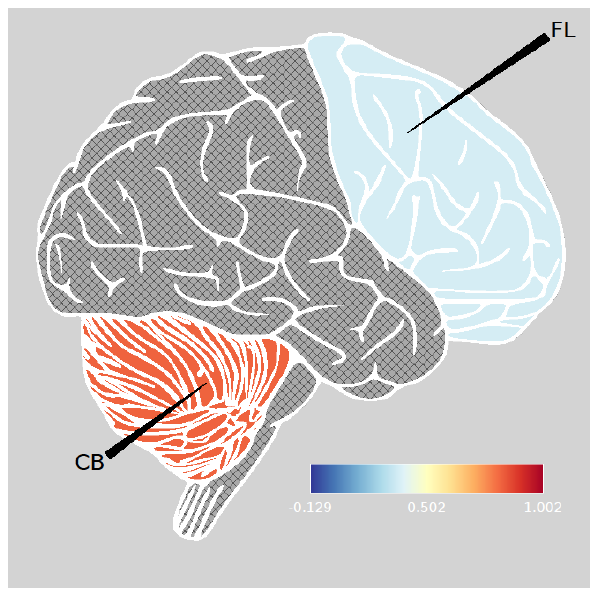

Figure S11: The brain regions indicating the difference in ZNF391 GReX between patients and controls with a color hue bar quantifying the degree of divergence between patients and controls; The brain regions correspond to the ones from human brain transcriptome (HBT) database

| 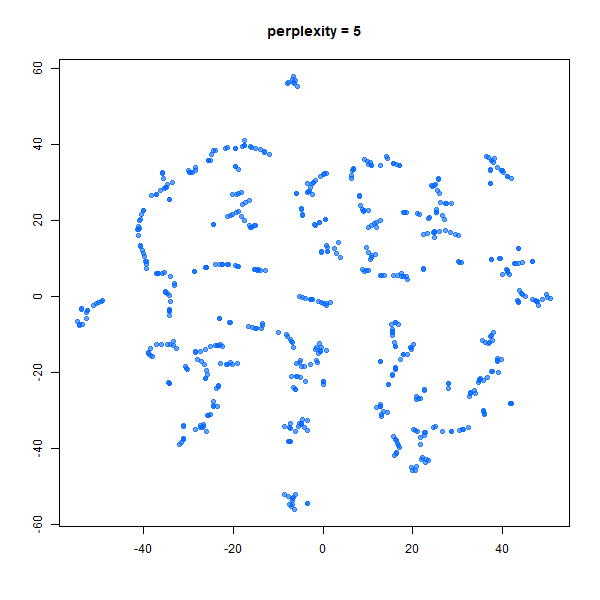(1) | 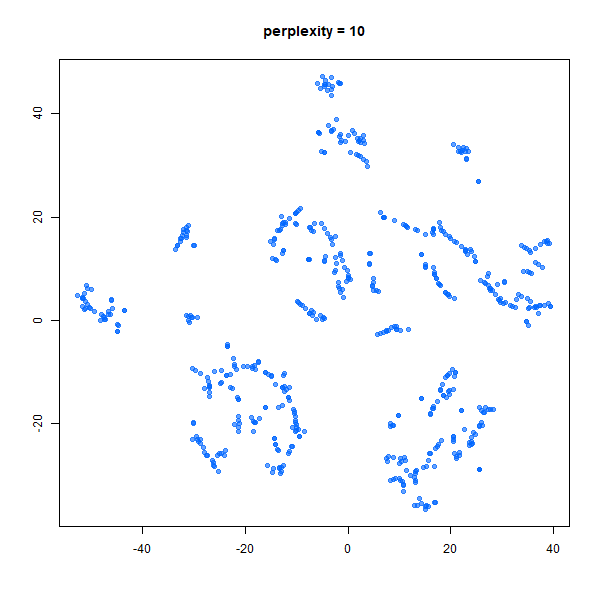  (2) | 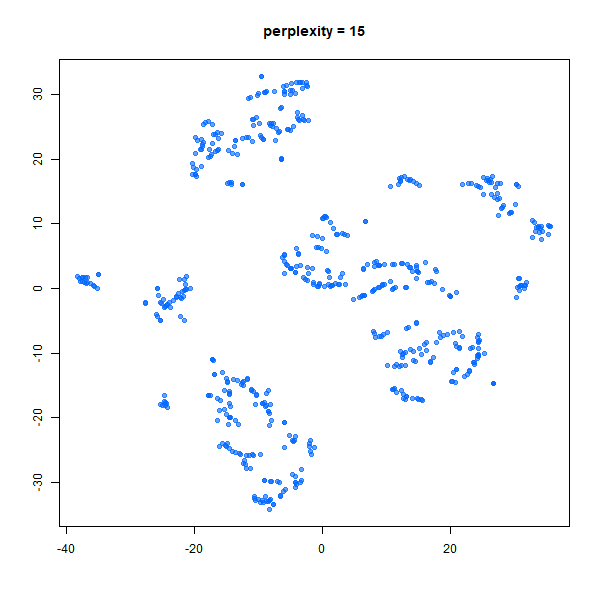  (3) | 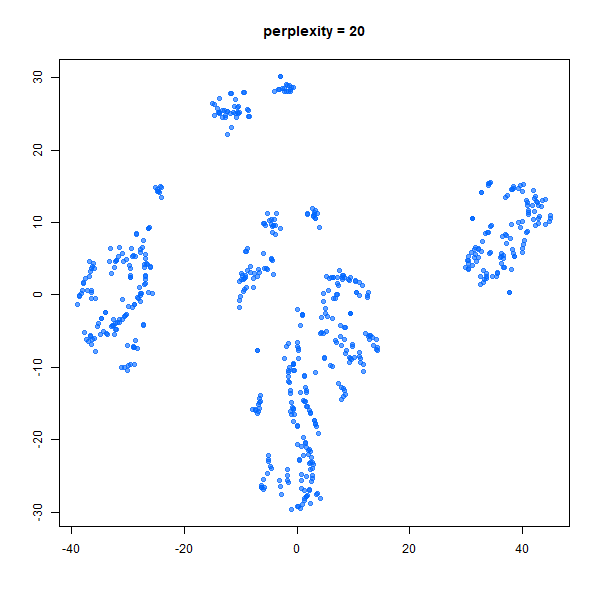  (4) | 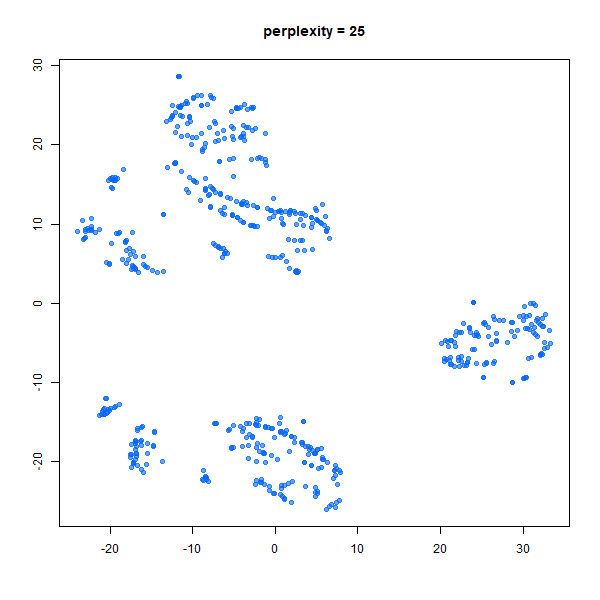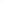(5) |
| --- | --- | --- | --- | --- |
| 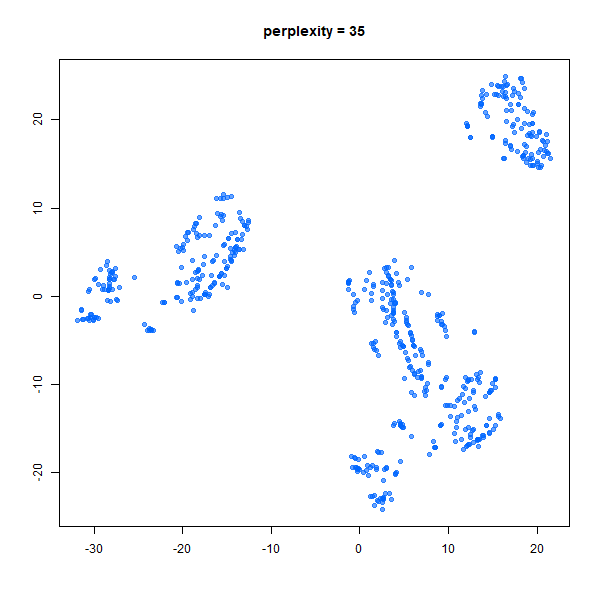  (6) | 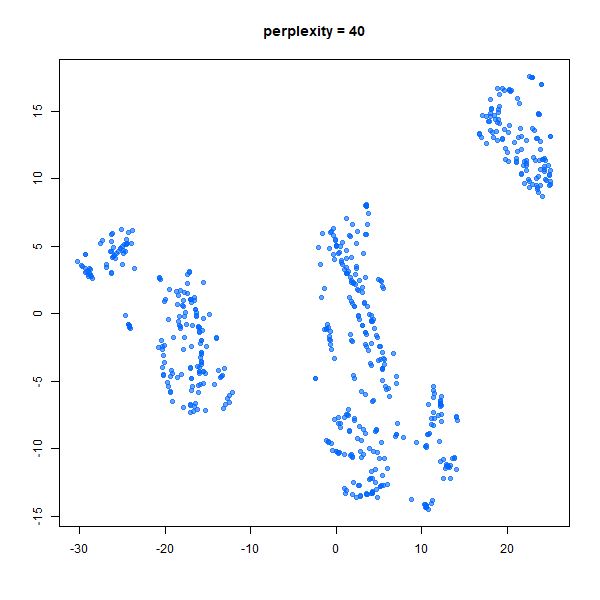  (7) | 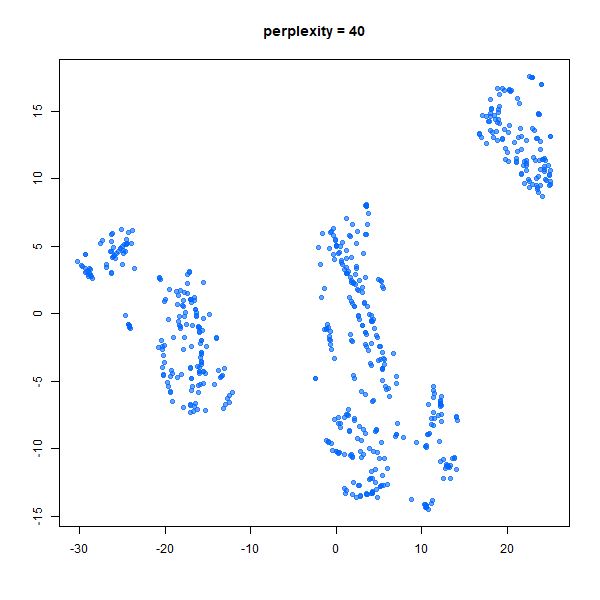  (8) | 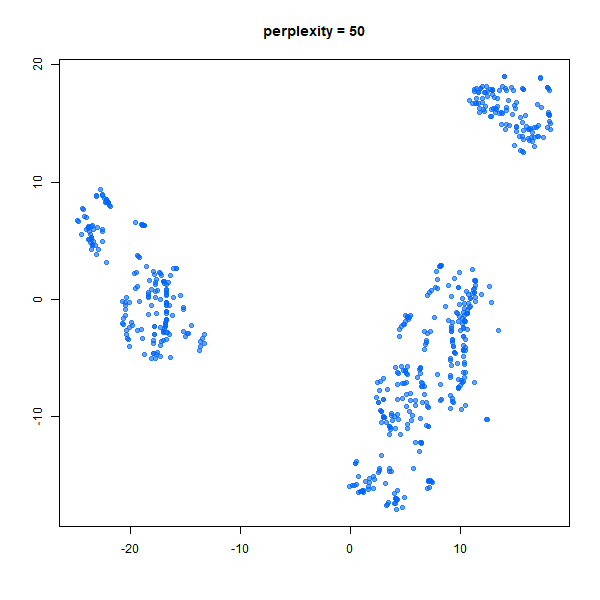  (9) | 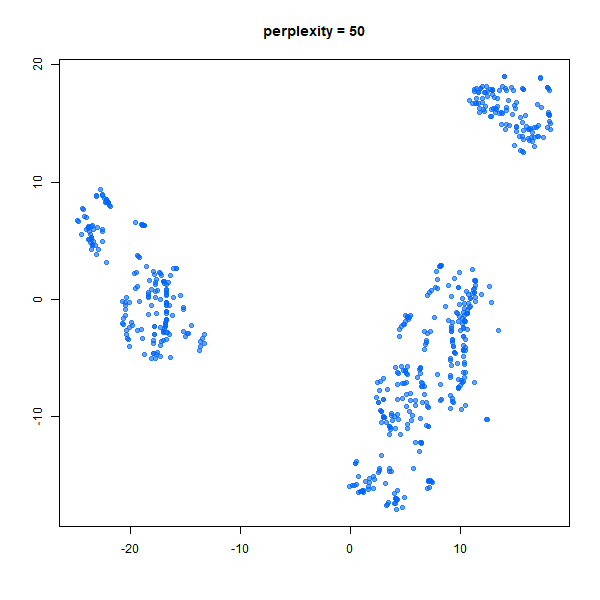  (10) |
| 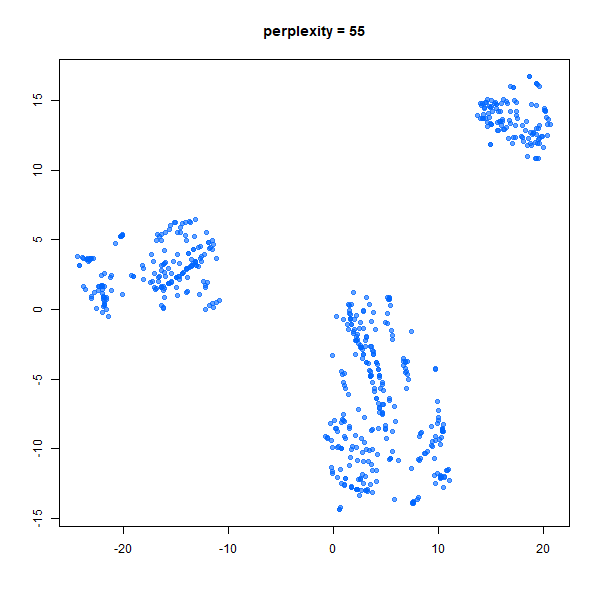  (11) | 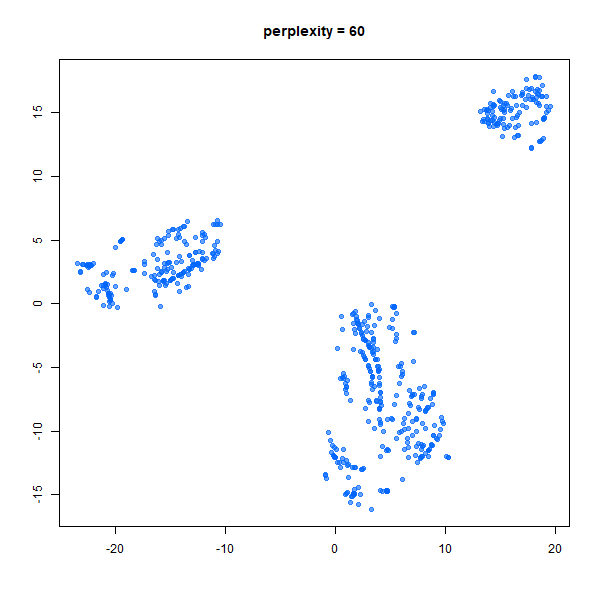  (12) | 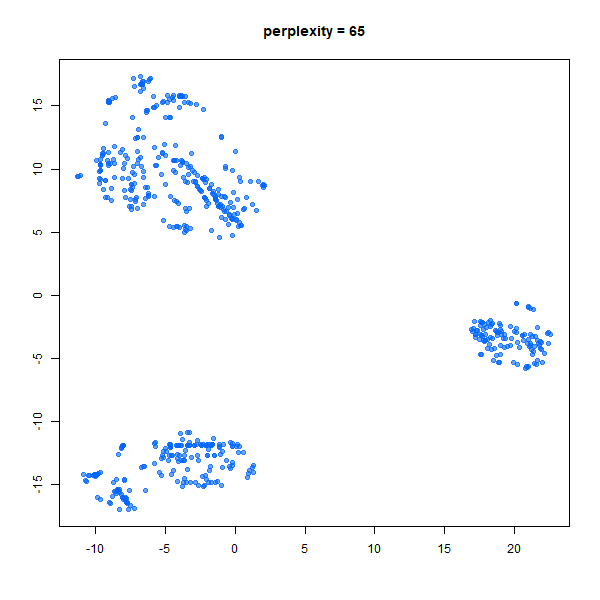  (13) | 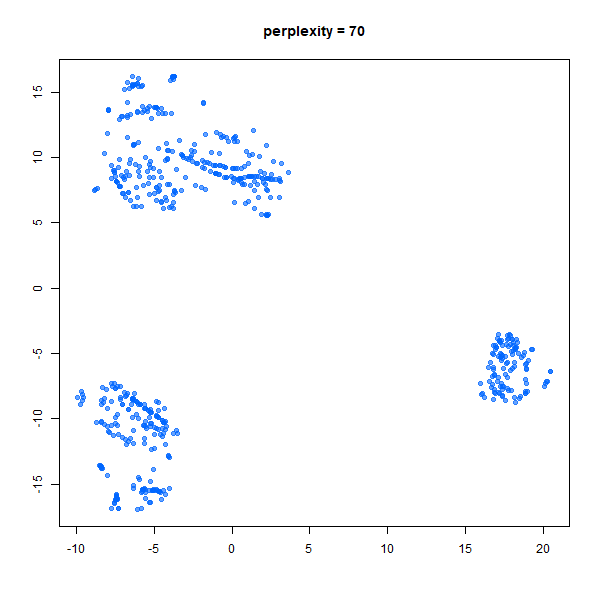  (14) | 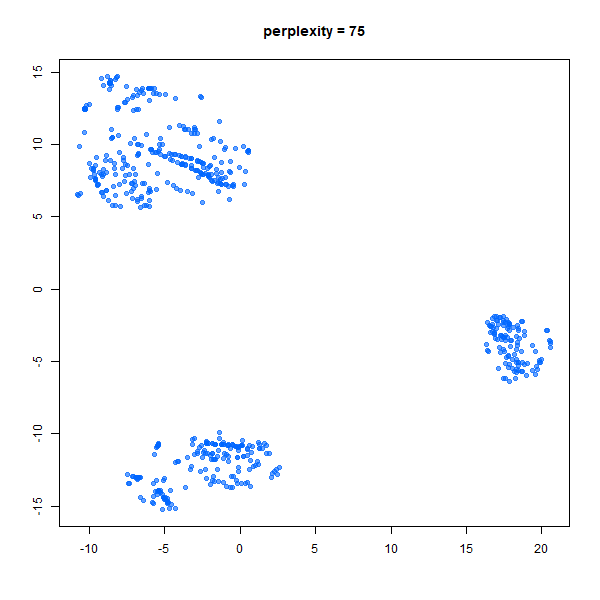  (15) |
| 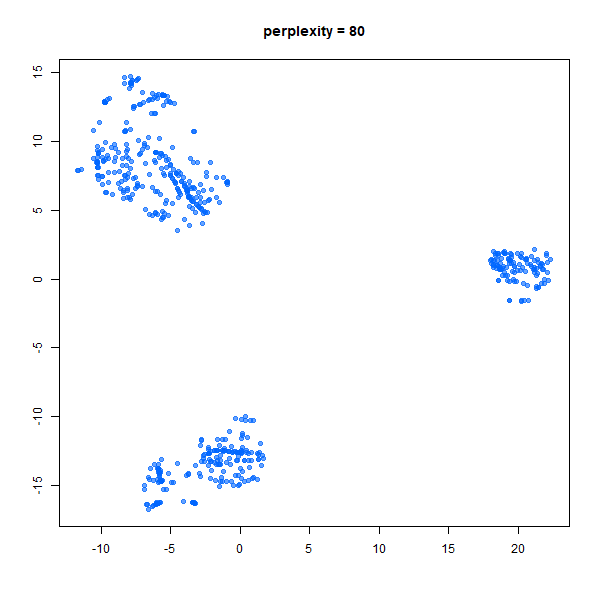  (16) | 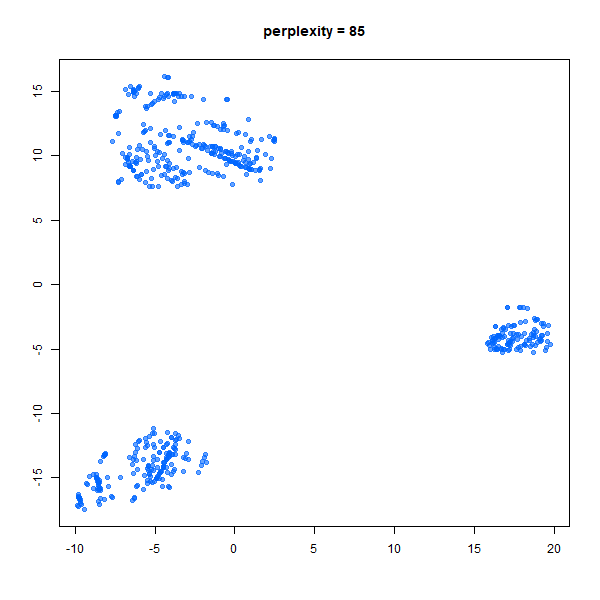  (17) | 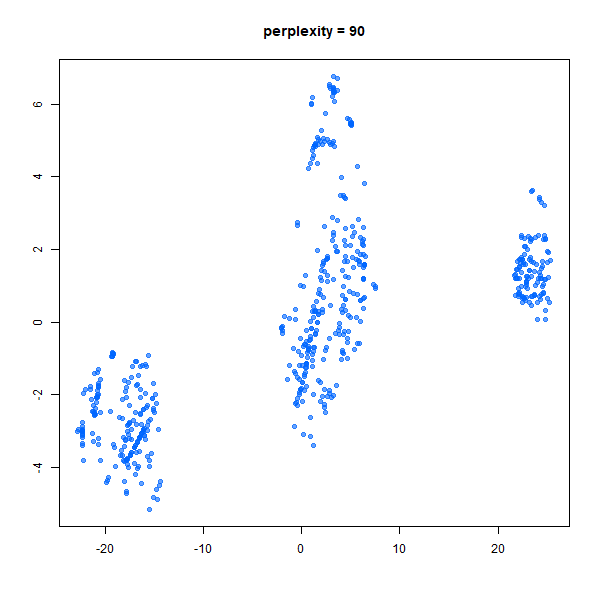  (18) | 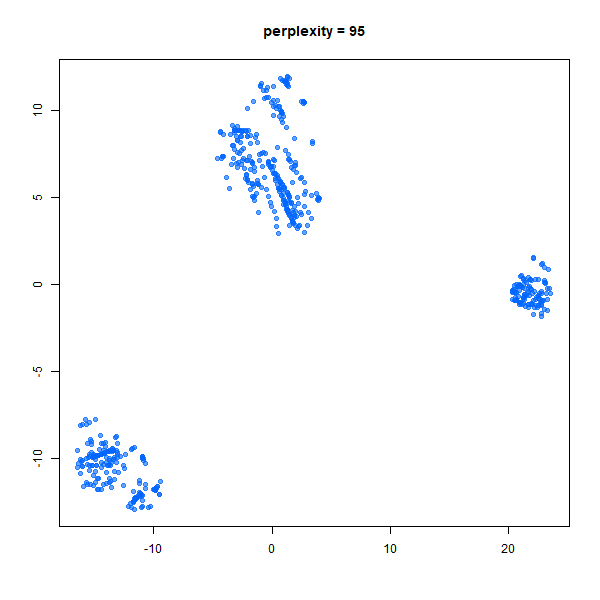  (19) | 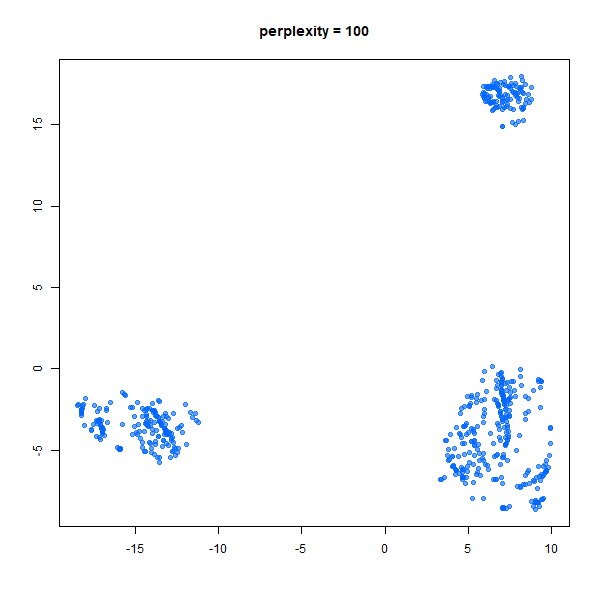  (20) |
| 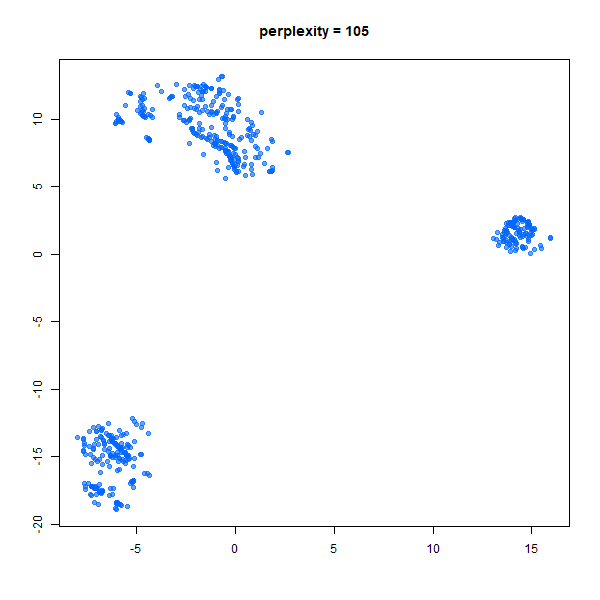  (21) | 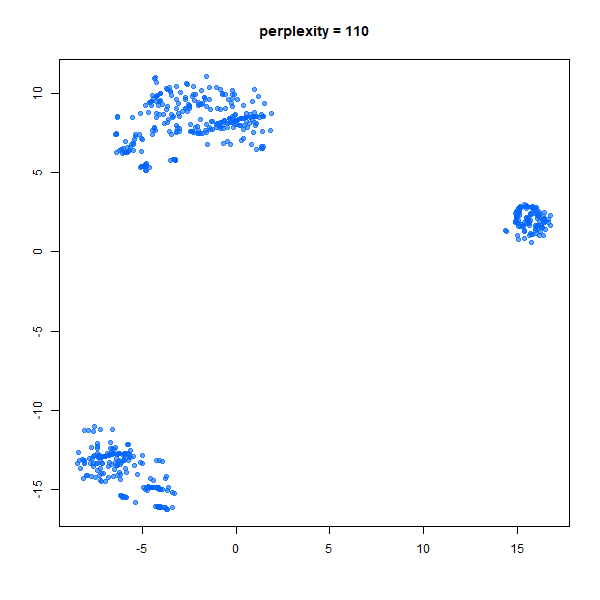  (22) | 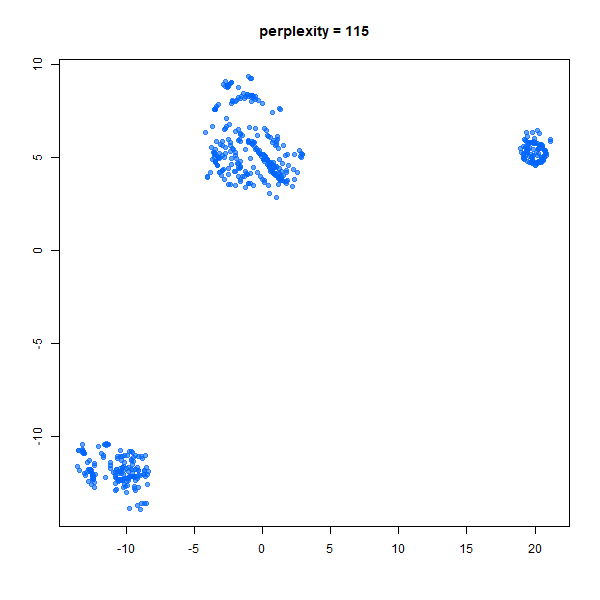  (23) | 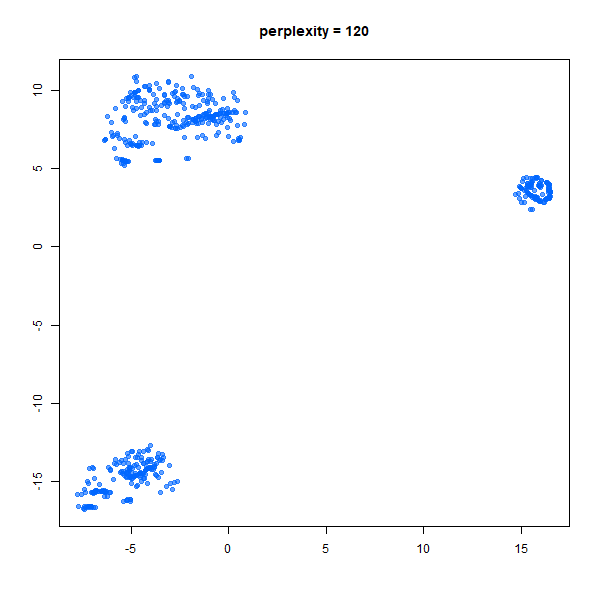  (24) | 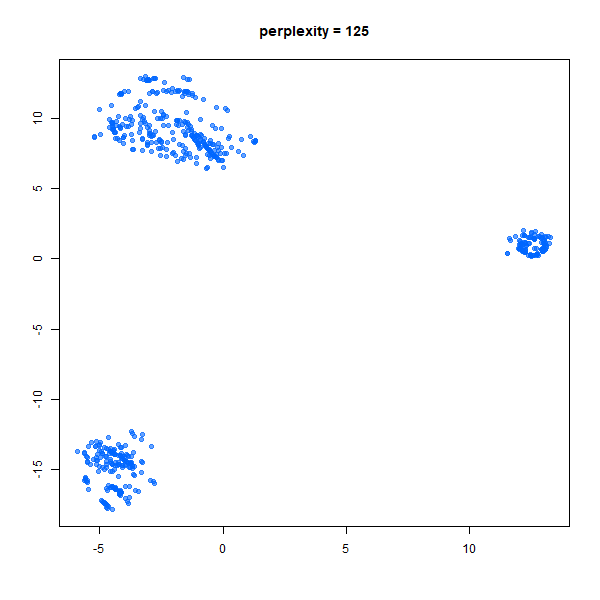  (25) |
| 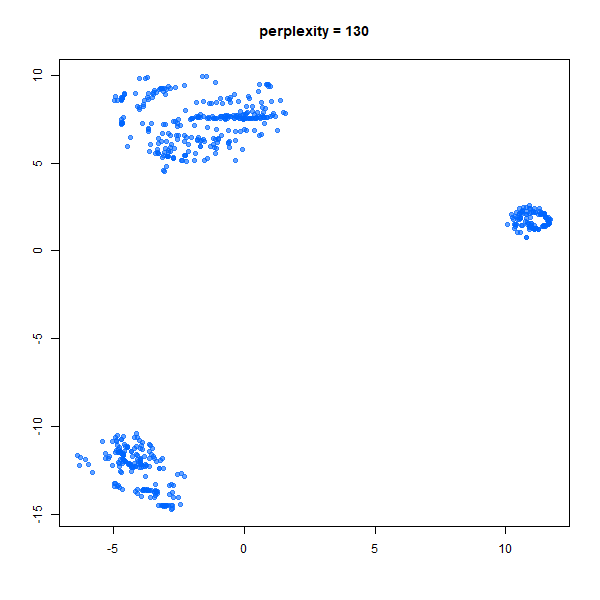  (26) | 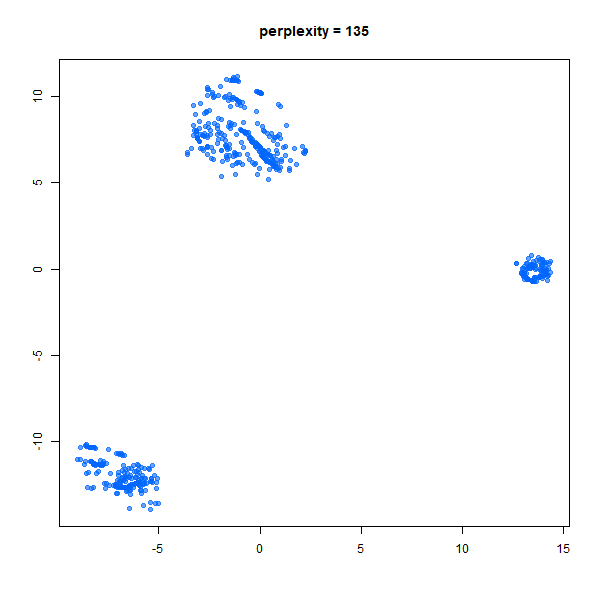  (27) | 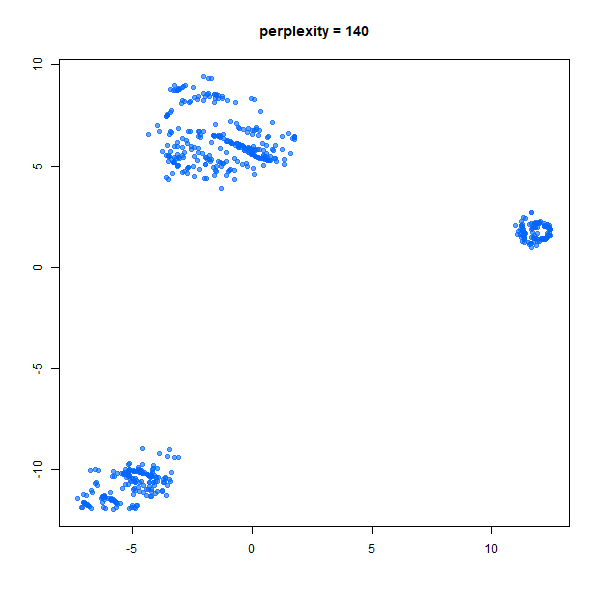  (28) | 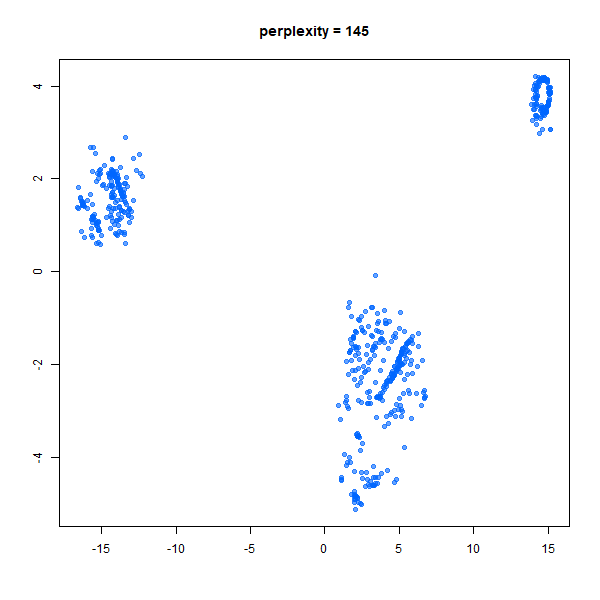  (29) | 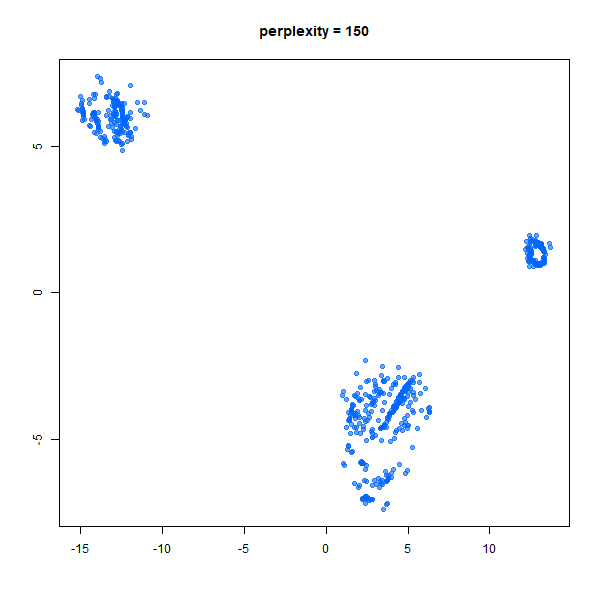  (30) |

Figure S13. Clustering pattern using different perplexity value in the t-SNE analysis

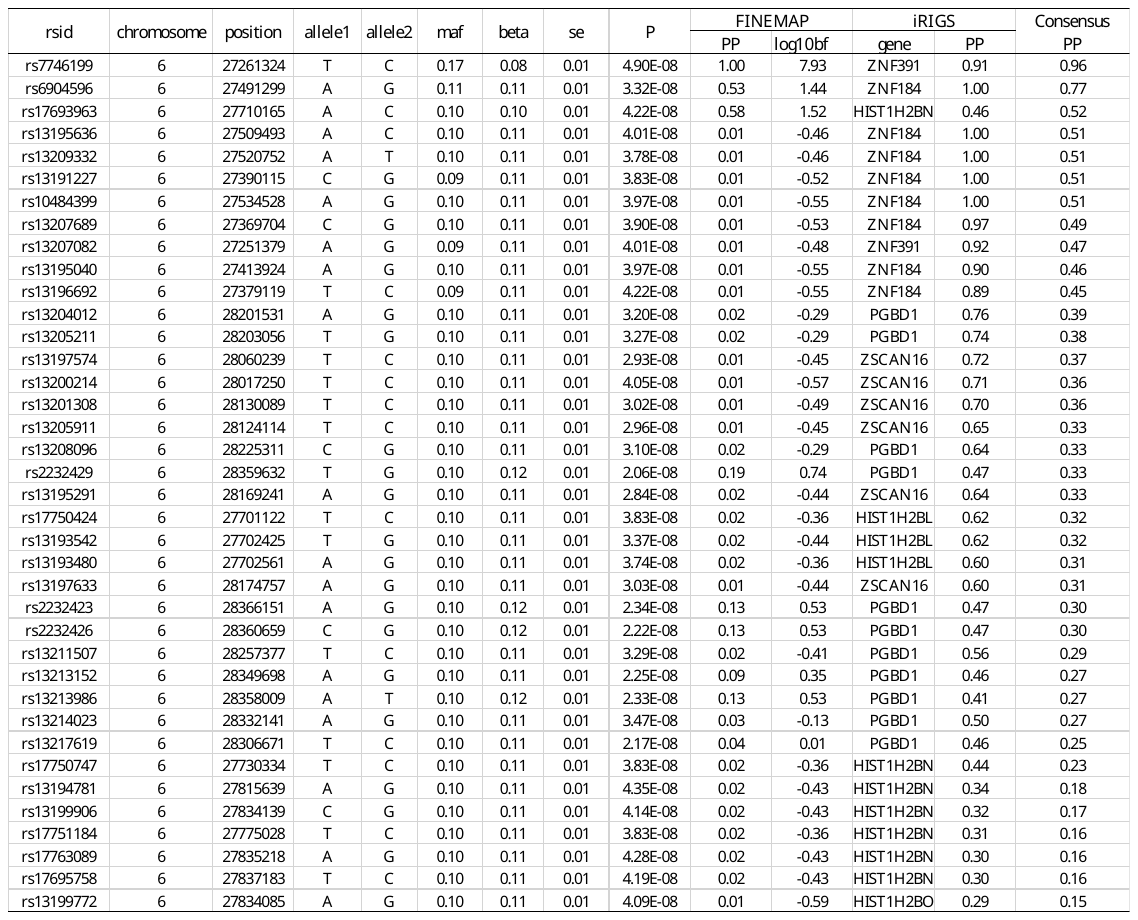

Table S2. Genome-wide scanning and fine-mapping results for the top loci (P < $5\times{10}^{-8}$) conferring risk to the common factor

| gene | genename | tissue | β | se | P |
| --- | --- | --- | --- | --- | --- |
| ENSG00000124613 | *ZNF391* | Amygdala | 0.05706 | 0.02428 | 0.0190 |
| ENSG00000124613 | *ZNF391* | Caudate_basal_ganglia | 0.08229 | 0.03201 | 0.0103 |
| ENSG00000124613 | *ZNF391* | Cerebellar_Hemisphere | 0.08975 | 0.03680 | 0.0149 |
| ENSG00000124613 | *ZNF391* | Cerebellum | 0.07710 | 0.03315 | 0.0202 |
| ENSG00000124613 | *ZNF391* | Frontal_Cortex | 0.04482 | 0.01546 | 0.00383 |
| ENSG00000124613 | *ZNF391* | Hypothalamus | 0.08561 | 0.03498 | 0.0146 |
| ENSG00000124613 | *ZNF391* | Substantia_nigra | 0.05668 | 0.02361 | 0.01656 |
| ENSG00000124613 | *ZNF391* | DLPFC | 0.014726 | 0.006169 | 0.0172 |

Table S3. The details of the comparison of ZNF391 GReX in the eight brain regions between patients and controls; β is the difference with patients as the reference group after controlling for three population principal components (PCs) and batch effect
